# Single-cell multiomic approaches define a gradual, spatially-regulated epigenetic and transcriptional transition from embryonic to adult neural stem cells

**DOI:** 10.64898/2026.02.06.704429

**Authors:** Beatrix S. Wang, Konstantina Karamboulas, Nareh Tahmasian, Daniel J. Dennis, David R. Kaplan, Freda D. Miller

## Abstract

Here, we ask how adult neural stem cells (NSCs) arise developmentally, focusing on murine cortical precursors that generate excitatory neurons embryonically and interneurons and glial cells postnatally. Using complementary single-cell spatial, transcriptomic, and epigenomic approaches, we show that postnatal NSC state acquisition involves a gradual transcriptional and epigenetic shift in the entire embryonic cortical precursor cell population and identify a distinct transition precursor state at E17/18 when both embryonic and the first postnatal progeny are being generated. Non-proliferative adult NSCs are also first seen at this transition timepoint, but they arise in a spatial domain distinct from that of the first postnatal progeny, indicating that NSC state acquisition is not a necessary prelude to the switch in cell genesis. These findings support a gradual epigenetically-continuous model for the transition from developing cortical precursors to NSCs and show that this is spatially separable from the transition to generating postnatal cell types.

## Introduction

A key question in mammalian stem cell biology involves the establishment of somatic tissue stem cell pools that play essential life-long roles in tissue maintenance and repair. Developmental perturbations in these pools can result in long-lasting adult impairments, and it is therefore essential to understand how they are initially established. One tissue where such developmental disruptions are particularly problematic is the mammalian brain, which contains two well-studied populations of neural stem cells (NSCs). One of these resides in the hippocampal dentate gyrus and generates postnatal neurons that are important for the flexibility of memory formation. The second population resides in the forebrain ventricular-subventricular zone (V-SVZ) and generates olfactory bulb interneurons and glial cells throughout life. Both of these NSC populations are well-studied with regard to their biological and functional roles,^1,2^ and a number of studies have defined the timeframe when these NSCs are generated developmentally.^3–7^ However, we still don’t know how an embryonic precursor transitions to an adult NSC, nor do we understand the molecular underpinnings of this important developmental process. Here, we have addressed these issues, focusing on murine forebrain NSCs.

Previous work has shown that postnatal forebrain NSCs have two distinct embryonic origins. The NSCs that reside along the lateral V-SVZ largely derive from embryonic ganglionic eminence (GE) radial precursors (RPs) that generate interneurons and oligodendrocytes throughout life. By contrast, NSCs that reside on the dorsal cortical side of the V-SVZ are generated by cortical RPs that make excitatory neurons embryonically, and then switch to making olfactory interneurons and glia postnatally.^8–10^ This switch in cell genesis occurs around the time of birth and is one of two major stem cell transitions that occur at this time.^4,11–13^ In the second transition, highly-active embryonic RPs give rise to slow-dividing, dormant/quiescent NSCs that persist throughout postnatal life.^14,15^ Further insights into the acquisition of this dormant state come from our previous work showing that (i) by postnatal day 6/7 (P6/7) cortical precursors display a transcriptionally-distinct NSC state and (ii) the transition to full NSC dormancy occurs over an extended timeframe commencing before P6/7 and ending by P21.^4^

These studies pinpoint the perinatal period as a time when cortical precursors become NSCs and start to make postnatal progeny. In this regard, one proposed explanation for these coincident events is the “set-aside” model, which posits that during mid-embryogenesis RPs destined to become NSCs are somehow selected and set aside as slowly-dividing precursors until they ultimately emerge as NSCs postnatally. In this model, there would be two populations of cortical RPs: those that generate excitatory neurons embryonically and those that are set-aside to become postnatal NSCs that make inhibitory neurons and glia. However, while several key studies indicate that the set-aside model is relevant for lateral GE-derived NSCs,^5,6^ a number of findings suggest it may not explain what happens in the cortex. First, Marcy and colleagues^7^ demonstrated that most dorsal cortical NSCs derive from precursors that are still proliferating around the time of birth. Second, clonal lineage tracing has demonstrated that at least some cortical precursors make excitatory neurons embryonically and glial cells and olfactory bulb interneurons postnatally.^5,16,17^ Finally, embryonic cortical RPs that make excitatory neurons can be induced to generate glia and interneurons when their local environment is changed by transplantation, exogenous ligands or manipulation of downstream signaling pathways.^12,13,18,19^ Thus, the set-aside model may not be applicable to the cortex. How then do embryonic cortical RPs become NSCs and what is the relationship between this state change and the genesis of postnatal cell types?

Here, we have used single-cell approaches to address these questions at the transcriptional, epigenetic, and spatial levels. We show that embryonic cortical RPs first start to generate postnatal cell types before birth, but that they largely do not acquire a non-proliferative NSC state until after birth. We show that acquisition of this NSC state involves a gradual transcriptional and epigenetic shift in the entire RP population and define a distinct transition precursor state that occurs at E17/18 when both embryonic and postnatal progeny are generated. Finally, we show that the first NSCs arise in a spatial domain that differs from the domain where the first postnatal progeny are generated. Thus, our findings support a continuous model for the transition from RPs to NSCs and show that this transition is spatially separable from the transition to generating postnatal cell types.

## Results

### At P2, cortical precursors have switched from generating excitatory neurons to making glia and interneurons

To determine when cortical precursors transition from making excitatory neurons to generating glia and olfactory interneurons, we defined both the precursors and their progeny using scRNA-seq from embryonic day 14 (E14) to postnatal day 6/7 (P6/7). We initially characterized the time immediately following birth since previous lineage tracing studies indicate that this is the time when the first cortically derived interneurons are generated.^4,20,21^ Specifically, we combined lineage tracing with scRNA-seq of the dorsal and lateral V-SVZ regions at postnatal day 2 (P2). To ensure we could distinguish cortically derived cells, we utilized mice carrying *Cre* knocked into the *Emx1* locus, and a floxed *Eyfp* allele in the *Rosa26* locus. We and others have used *Emx1Cre;R26Eyfpfl/fl* mice (from hereon called *Emx1-Eyfp* mice) to specifically tag cortically-derived cells with *Eyfp* expression.^4,22,23^ We dissected the dorsal/lateral V-SVZ from these mice at P2 and sequenced single cells using the 10x Genomics Chromium platform. We analyzed the resultant transcriptomes using our previously described pipeline.^4,24–29^ After filtering the datasets for doublets and low-quality cells with, for example, few expressed genes or high (>12%) mitochondrial DNA content, we used genes with high variance to compute principal components as inputs for projecting cells in two-dimensions using UMAPs. Clustering was performed using a shared nearest neighbors-cliq (SNN-cliq)-inspired approach built into the Seurat R package at a range of resolutions.^30^

We identified all predicted cell types within this dataset, including neural stem cells (NSCs) and their postnatal progeny, transit-amplifying precursors (TAPs), neuroblasts, oligodendrocyte precursor cells (OPCs), and newborn astrocytes as well as newborn excitatory neurons, striatal interneurons, and microglia (Figure S1A). We defined postnatal NSCs as cells that expressed the NSC marker mRNAs *Aldoc*, *Slc1a3/Glast*, *Tfap2c*, *Vnn1*, and *Veph1* but that expressed low levels of astrocyte mRNAs like *Agt* and *Htra1*. We confirmed this assignment using a previously defined gene signature^4^ that distinguishes postnatal NSCs from astrocytes (Figure S1B). We defined TAPs as proliferative cells expressing well-characterized marker mRNAs such as *Ascl1* and *Egfr*, as well as other TAP genes we previously defined,^4,28^ but not expressing OPC, astrocyte or NSC genes. We defined intermediate progenitors (IPs) for the excitatory neuron lineage as cells expressing the well-characterized IP marker mRNAs *Eomes* and *Insm1* but not neuronal mRNAs such as *Neurod1* and *Neurod6*. We considered these IPs to be early in the neurogenic lineage and therefore a reflection of ongoing excitatory neurogenesis; as such, we termed them early IPs. Since this P2 dataset did not include any early IPs (Figure S1A), this indicates that excitatory neurogenesis is largely complete by P2. Finally, we identified cells that expressed early neuronal mRNAs such as *Neurod1* and *Neurod6*, as well as the IP mRNA *Eomes*; these are newborn neurons that have likely just been generated from IPs (see Yuzwa et al.^29^) and we have therefore termed them late IPs/newborn neurons.

To increase the power of our analysis, we merged these transcriptomes with our previously-published P2 *Emx1-Eyfp* mouse V-SVZ dataset (Borrett et al.,^4^ GEO: GSE152281) after reanalyzing it in the same way (Figure S1C). We used two iterations of the Harmony batch correction approach to ensure integration of the two datasets.^31^ Marker gene analysis of the merged dataset identified the same cell types as in the individual datasets (Figures 1A and 1B). Analysis of the cell cycle using Cyclone^32^ (Figure 1C) predicted that many of the TAPs were proliferating, as were some OPCs and astrocytes. Neither NSCs nor late IPs/newborn neurons were predicted to be proliferative.

**Figure 1.**
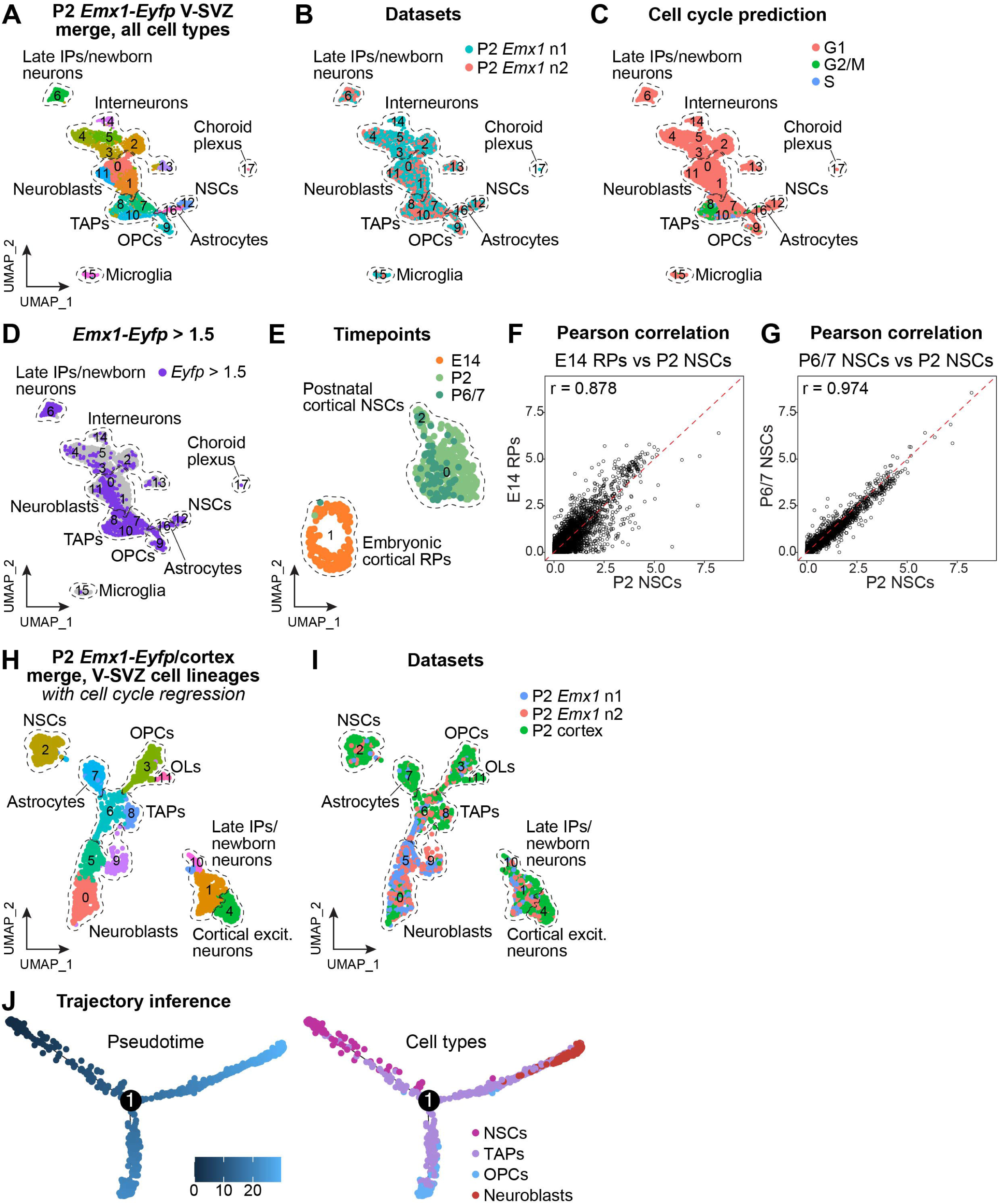
Single-cell RNA-seq shows that by P2 cortical precursors have acquired an NSC transcriptional state. **(Also see Figure S1).(A-D)** The P2 V-SVZ from *Emx1-Eyfp* mice was analyzed by two independent scRNA-seq runs (see Figures S1A and S1C), and the resultant datasets were merged and batch-corrected with two iterations of Harmony. **(A, B)** UMAP projections of the merged P2 dataset showing (A) transcriptionally distinct color-coded clusters, annotated for cell types (indicated by hatched lines) and (B) datasets of origin, colored as per the adjacent color key. **(C)** UMAP of the dataset in (A), analyzed by Cyclone to predict cell cycle status, with different cell cycle phases annotated as per the adjacent color key. **(D)** UMAP of the dataset in (A) overlaid for expression of *Emx1-Eyfp* mRNA. Normalized expression levels were thresholded so that cells with levels ≥ 1.5 are shown as purple and other cells as grey. **(E)** UMAP of merged E14, P2, and P6/7 *Emx1-Eyfp*-expressing RP/NSCs, color-coded for timepoint as per the adjacent key. Transcriptionally-distinct clusters are numbered and outlined by hatched lines. The P2 NSCs are from the dataset in Figure S1D (cluster 12) and the E14 RPs and P6/7 NSCs are shown in Figures S1E and S1G. **(F, G)** Pearson correlation plots comparing averaged gene expression of *Emx1-Eyfp* P2 NSCs with *Emx1-Eyfp* E14 RPs and *Emx1-Eyfp* P6/7 NSCs, all shown in (G). Each dot represents an individual gene, r values indicate the Pearson correlation coefficient, and red hatched lines indicate r values of 1. **(H, I)** UMAPs of P2 cortex V-SVZ lineage neural cells, generated by merger of the datasets in Figures S1D and S1H following cell cycle regression and two iterations of Harmony, showing (E) transcriptionally-distinct color-coded clusters, annotated for cell types (indicated by hatched lines) and (F) datasets of origin, colored as per the adjacent color key. **(J)** Cell cycle regressed Monocle2-based pseudotime ordering of P2 NSC, TAP, OPC, and neuroblast transcriptomes, all extracted from the dataset in (H). Cells are colored based on pseudotime scores (left) or cell type annotations (right), as per the relevant keys. **Abbreviations:** Excit., excitatory. IPs, intermediate progenitors. NSCs, neural stem cells. OLs, oligodendrocytes. OPCs, oligodendrocyte precursor cells. RPs, radial precursors. TAPs, transit-amplifying precursors.

Analysis of *Eyfp* expression within this merged dataset (Figure 1D) showed that approximately 80% of the late IPs/newborn excitatory neurons expressed *Eyfp* mRNA while cortical/striatal interneurons displayed only low ambient RNA levels of *Eyfp* expression, confirming the efficacy and specificity of the lineage tracing. Many TAPs, neuroblasts, OPCs, and astrocytes were also *Eyfp*-positive, indicating they were generated by cortical precursors. Since there are no early IPs within this merged dataset (Figure 1A), then this indicates that by P2 cortical precursors have largely ceased generating excitatory neurons and are instead making glia and olfactory bulb interneurons.

### By P2, cortical precursors have acquired an NSC transcriptional state

To confirm that P2 cortical precursors had acquired an NSC-like state and to better define this state, we compared them transcriptionally to more mature cortical NSCs. To ensure we only analyzed cortical stem cells, we subsetted out *Eyfp*-positive neural cells from the merged P2 dataset (Figure S1D). We compared these to transcriptomes of *Emx1-Eyfp* lineage-traced E14 cortical radial precursors (RPs) and P6/7 NSCs that we previously characterized and published (Borrett et al.,^4^ GEO: GSE152281), subsetting out the cortically-derived *Eyfp*-positive cortical precursors in both cases (Figures S1E-S1G). In the resultant merged precursor dataset (Figure 1E), the P2 NSCs co-clustered with the P6/7 NSCs, while the E14 RPs were completely distinct. Consistent with this, comparison of the RP/NSCs from the three timepoints using Pearson correlation of averaged gene expression (Figures 1F and 1G) showed that the P2 cells were highly similar to the P6/7 NSCs, with r = 0.974, but not to the E14 cortical RPs, with r = 0.878.

These findings indicate that, by P2, cortical precursors have acquired a postnatal NSC transcriptional state and have switched to generating their adult progeny. To extend these conclusions, we performed Monocle-based trajectory analysis on P2 cortical NSCs and their immediate progeny. We enhanced the power of this analysis by including transcriptomes from our published P2 cortex scRNA-seq dataset where the cortex was carefully dissected prior to analysis (Dennis et al.,^25^ GEO: GSE255405) (Figure S1H). We subsetted the cortical V-SVZ neural lineage transcriptomes from this dataset and merged them with the P2 *Emx1-Eyfp*-positive V-SVZ neural transcriptomes (those in Figure S1D). We performed batch correction and cell cycle regression on this merged dataset (Figures 1H-1I) to ensure results were not driven by cell cycle status. Monocle-based trajectory analysis of the NSCs, TAPs, OPCs and neuroblasts from this dataset (Figure 1J) predicted a trajectory with NSCs at one end, followed by TAPs and then a bifurcation giving rise to OPCs and neuroblasts, a trajectory similar to that seen with adult NSCs.

### At E17/18, cortical precursors are in a unique RP to NSC transition state and start to generate the olfactory interneuron lineage

These findings indicate that the transition from cortical RPs to NSCs occurs between E14 and P2, consistent with previous work.^4,7^ To more precisely define the transition, we performed scRNA-seq on E18 cortical tissue. Analysis of the resultant dataset identified all major cortical cell types including excitatory neurons, cortical interneurons, IPs, OPCs, microglia, and cortical precursor cells (RP/NSCs) (Figure S2A). We also identified TAPs and olfactory bulb neuroblasts in this E18 dataset, suggesting that cortical precursors start generating postnatal progeny prior to birth.

To explore this latter idea definitively, we merged the E18 cortical neural cells with E17 cortically-derived *Emx1-Eyfp*-positive neural cells from our previously published E17 *Emx1-Eyfp* V-SVZ dataset (Borrett et al.,^4^ GEO: GSE152281) and performed a single iteration of Harmony to account for batch effects (Figure S2B). Analysis of this merged E17/18 cortical dataset (Figures 2A-2C) identified *Eyfp*-positive TAPs, olfactory bulb neuroblasts, and OPCs, confirming that cortical precursors generate postnatal progeny at E17/18. This merged dataset also included early IPs expressing *Eomes*, *Insm1*, and *Pax6*, but not *Neurod1* or *Neurod6* (Figures 2A and S2C), indicating that excitatory neurogenesis was also still ongoing. Moreover, Cyclone analysis showed that at this age some RP/NSCs were still proliferative (Figure S2D). Thus, unlike at P2, at E17/18 RP/NSCs are still proliferating, and as a population they are generating both the excitatory neuron lineage, as indicated by the presence of early IPs, and the olfactory interneuron lineage, as indicated by the presence of cortically-derived TAPs.

**Figure 2.**
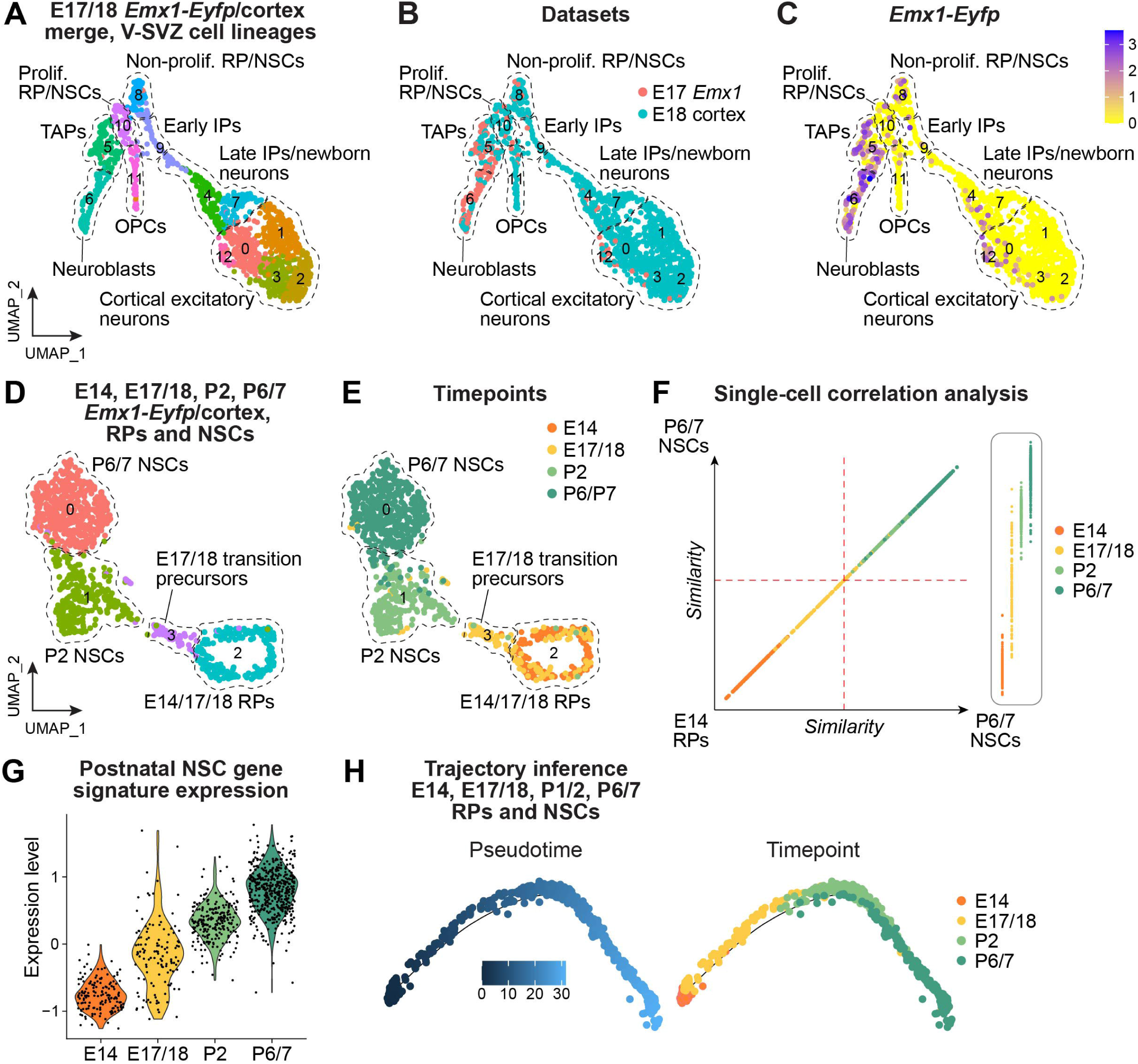
scRNA-seq demonstrates that cortical precursors undergo a gradual transcriptional shift from RP to NSC states between E14 and P2. **(Also see Figure S2).(A-C)** UMAPs of V-SVZ cell lineage transcriptomes from a merger between the E18 cortex dataset (Figure S2A) and cortically-derived *Emx1-Eyfp*-positive transcriptomes from our previously published E17 *Emx1-Eyfp* V-SVZ dataset (Borrett et al.,^4^ GEO: GSE152281; see Figure S2B) after one round of batch correction. Shown are **(A)** transcriptionally-distinct color-coded clusters, annotated for cell types (indicated using hatched lines), (B) datasets of origin, color-coded as per the adjacent key, and (C) *Emx1-Eyfp* expression levels, color-coded as per the adjacent key. Note that only one of the two merged datasets is lineage-traced, and the unlabelled cells in **(C)** come from the other dataset. **(D, E)** UMAPs of merged, batch-corrected E14, E17/18, P2, and P6/7 cortical precursor transcriptomes, showing **(D)** transcriptionally-distinct, color-coded clusters (numbered and outlined with hatched lines), and (E) timepoints of origin, as per the adjacent color key. E14 transcriptomes come from Figure S1E, E17/18 transcriptomes from the dataset in (A, clusters 8 and 10), P2 transcriptomes from Figure 1H (cluster 2), and P6/7 transcriptomes from Figure S2E (cluster 3). Two iterations of Harmony were performed. **(F)** Individual RP and NSC transcriptomes from the dataset in (D) were Pearson correlated against averaged E14 RP and P6/7 NSC transcriptomes. Shown is a scatter plot in which individual cells are plotted based on similarity to E14 RPs vs P6/7 NSCs along both the x- and y-axes. Cells are color-coded based on timepoint. The inset on the right shows the location of transcriptomes from each individual timepoint along the y-axis. **(G)** Violin plot showing expression of a previously-defined postnatal NSC gene signature (from Borrett et al.^4^ with slight modifications) in E14, E17/18, P2, and P6/7 cortical precursors from the dataset in (D), separated by timepoint. Each dot represents gene signature levels in an individual cell. **(H)** Cell cycle regressed Monocle2-based pseudotime ordering of the E14, E17/18, P2, and P6/7 cortical precursors from the dataset in (D). Cells are colored based on pseudotime scores (left) or timepoint (right) as per the adjacent keys. **Abbreviations:** IPs, intermediate progenitors. NSCs, neural stem cells. OPCs, oligodendrocyte precursor cells. Prolif., proliferative. RPs, radial precursors. TAPs, transit-amplifying precursors.

We next asked whether the E17/18 cortical precursors were more like RPs or more like NSCs by comparing them to cortical stem cells from E14 to P6/7. To do this, we merged the E17/18 cortical precursors (clusters 8 and 10 in Figure 2A) with E14 cortical RPs (Figure S1E) and P2 cortical NSCs (cluster 2 from Figure 1H). We also included P6/7 cortical NSCs subsetted from a merger of the P6/7 *Emx1-Eyfp* datasets and our published P7 dorsal cortex scRNA-seq dataset (Dennis et al.,^25^ GEO: GSE255405) (cluster 3 from the merged dataset in Figures S2E-S2F). To reduce batch effects, two iterations of Harmony were performed. The resultant transcriptional comparison (Figures 2D-2E) showed that some E17/18 precursors clustered with E14 RPs, a few with P2 NSCs, but that many clustered as a separate, transcriptionally-distinct group (cluster 3). We have termed these distinct E17/18 cells transition precursors.

We further characterized this stem cell dataset using four approaches. First, we performed single-cell correlation analysis, comparing each cortical precursor transcriptome to the averaged gene expression for E14 cortical RPs versus P6/7 cortical NSCs. This analysis (Figure 2F) demonstrated a gradual transcriptional transition from E14 to P6/7, with the E17/18 precursors showing almost equal similarity to the RPs and NSCs. Second, we characterized the onset of postnatal NSC gene expression using a gene signature we had previously defined, and modified slightly (Borrett et al.,^4^ see Experimental Methods). This NSC transcriptional signature was robustly expressed by P2 and P6/7 NSCs and was lower in E14 RPs (Figure 2G). Notably, the E17/18 cells were heterogeneous, with most displaying expression values intermediate between RPs and NSCs. Consistent with this, violin plots showed that postnatal NSC genes like *Gfap* and *Aqp4* were just starting to increase in the precursors at E17/18 (Figure S2G). Third, we performed Monocle-based trajectory analysis on the E14 to P6/7 cortical precursors after regressing out cell cycle genes to ensure it was not driven by proliferation status. This analysis (Figure 2H) identified a single trajectory consistent with a gradual transition from RPs to postnatal NSCs. Specifically, E14 RPs were at one end, followed by the E17/18 transition precursors and then the P2 NSCs and P6/7 NSCs, with substantive overlap between the two postnatal NSC populations. Notably, the majority of E17/18 transition precursors occupied a distinct trajectory location. Thus, cortical precursors undergo a gradual population shift from RPs to NSCs between E14 and P2, with a transition midpoint around E17/18 when they express both RP and NSC genes. Moreover, as a population they generate both early IPs for the excitatory neuron lineage and TAPs for the olfactory interneuron lineage.

### ***S***ingle-cell multiomics defines the transcriptional states and chromatin accessibility of cortical precursors from E14 to P7

We asked whether the gradual transcriptional transition from RPs to NSCs was driven at the chromatin level by acquiring single-nucleus RNA-seq (snRNA-seq) and single-nucleus assay for transposase accessible chromatin-seq (snATAC-seq) from the same single nuclei using the 10x Genomics platform. We analyzed cortical cells at E14 (prior to the transition to an NSC state), at E17 (during the transition), at P2 (after the transition), and at P6/7 (at which point dormant postnatal NSCs are well-established).^4^ The transcriptomic data was processed through our Seurat-based pipeline as for the scRNA-seq datasets, while the epigenomic data was processed using a separate pipeline based on the R package ArchR.^33^ After filtering out low quality cells based on metrics such as fragment count and transcription start site enrichment, we performed dimensionality reduction using an iterative latent semantic indexing approach. We clustered the data across a range of resolutions using Seurat’s graph-based clustering approach^30^ and visualized the datasets in two-dimensional space using UMAP embeddings.

We first analyzed the E14 dorsal cortex and obtained 4,780 paired transcriptomes and epigenomes. We clustered the dataset based on the transcriptional data and identified all of the expected cell types, including excitatory neurons, cortical interneurons, microglia, RPs, and IPs (Figure S3A). We subsetted out the V-SVZ lineage neural cells and performed Cyclone-based cell cycle prediction. This analysis defined proliferative and non-proliferative RPs, as well as proliferative and non-proliferative IPs (Figures 3A and S3B). We then reclustered the E14 cortical V-SVZ cells based upon chromatin accessibility. Marker gene accessibility identified the same cell types as with the transcriptome-based clustering (Figure 3B). We compared the transcriptomic versus chromatin accessibility-based cell type identification by overlaying bar codes for a given epigenetically-defined cell type on the transcriptome-based UMAP (Figure 3C). This analysis confirmed close correspondence between the two approaches but also defined two differences. First, proliferative and non-proliferative cells were not distinguishable using the chromatin accessibility data (for example, see IPs in Figures 3A-3C). Second, epigenome-defined changes in cell type sometimes preceded those observed transcriptionally. For example, some cells transcriptionally-defined as RPs clustered instead with early IPs in the chromatin-based clustering, presumably because genes in differentiating cells became accessible prior to detectable mRNA expression.

**Figure 3.**
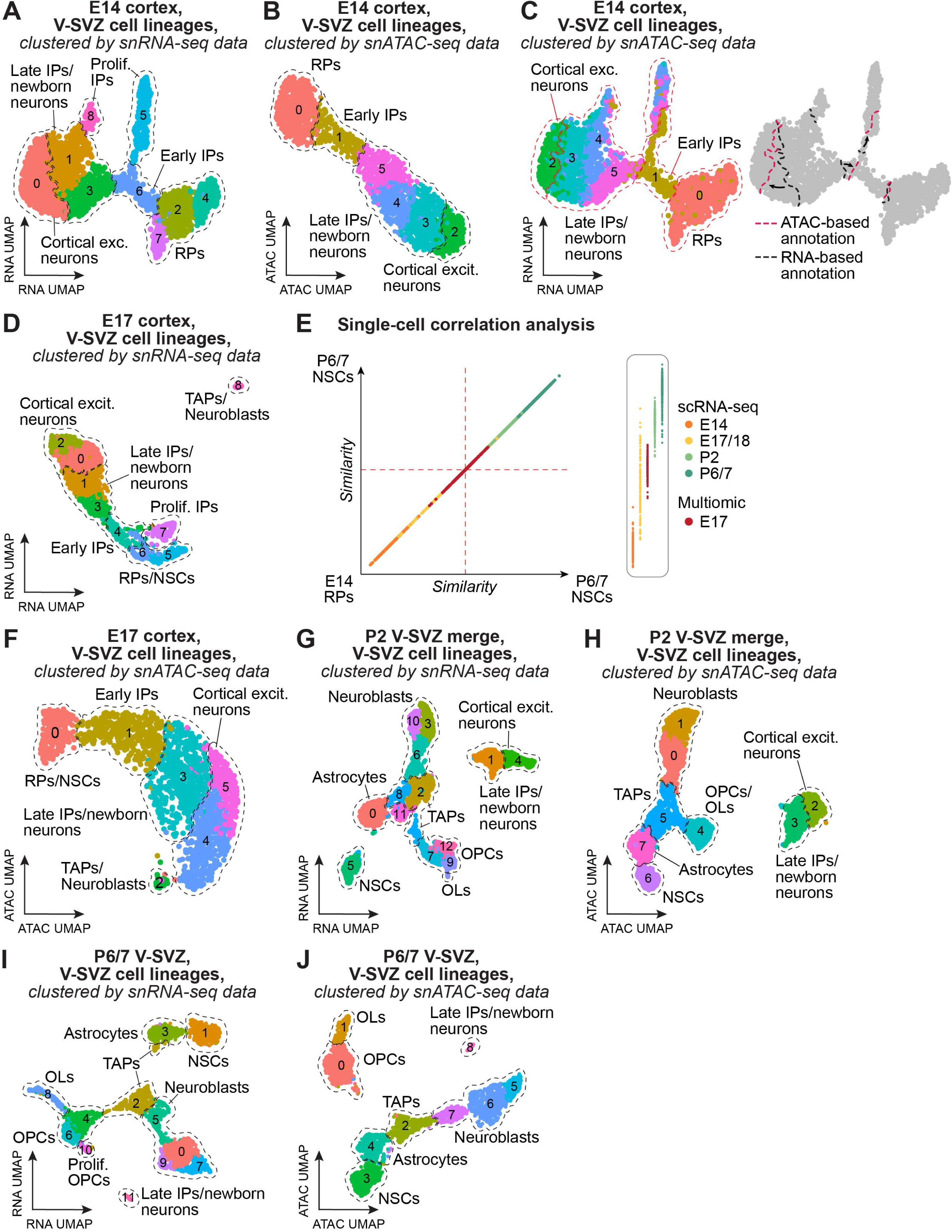
Combined single nucleus RNA-seq and ATAC-seq multiomic analysis of cortical precursors from E14 to P6/7. **(Also see Figure S3).(A)** UMAP of the E14 cortex V-SVZ cell lineage (subsetted from the multiomic dataset in Figure S3A), clustered on the basis of the transcriptional data. Distinct clusters are colored and numbered and cell types are annotated and outlined based upon marker gene expression. **(B)** UMAP of the same E14 cortex V-SVZ cell lineage multiomic dataset shown in (A), clustered on the basis of the snATAC-seq chromatin accessibility data. Colors and numbers represent epigenetically-distinct cell clusters, and cell types are annotated and outlined based upon marker gene chromatin accessibility analysis. **(C)** UMAPs of the multiomic dataset from (A, B), comparing cell type assignments based on the transcriptomic versus epigenomic data. The UMAP plot on the left is from the transcriptome-based clustering (as in A), but the cell clusters are colored and annotated based on their assignments in the epigenomic clustering in (B). The UMAP on the right is the same as on the left but in this case borders of the annotated cell types are shown based on the transcriptional clustering versus epigenomic clustering are indicated as black and red lines, respectively. **(D)** UMAP of the E17 cortex V-SVZ neural cell lineage (subsetted from the multiomic dataset in Figure S3C), clustered on the basis of the transcriptional data. Distinct clusters are colored and numbered and cell types are annotated and outlined based upon marker gene expression. **(E)** Individual E17 cortical RP/NSC transcriptomes from the dataset in (D) were Pearson correlated against averaged E14 RP and P6/7 NSC scRNA-seq transcriptomes from the analysis shown in Figure 2F. Also included for comparison are the individual E14 to P6/7 RP/NSCs from the dataset in Figure 2D. Individual transcriptomes are plotted based on similarity to scRNA-seq-defined transcriptomes of E14 cortical RPs versus P6/7 cortical NSCs along both the x- and y-axes. The inset on the right shows the location of transcriptomes from each individual timepoint along the y-axis. Cortical precursors from different timepoints are color-coded as per the adjacent key. **(F)** UMAP of the same E17 cortex V-SVZ neural cell lineage multiomic dataset shown in (D), clustered on the basis of the snATAC-seq chromatin accessibility data. Colors and numbers represent epigenetically-distinct cell clusters, and cell type annotations (indicated by the hatched lines) are based upon marker gene chromatin accessibility analysis. **(G)** UMAP showing the P2 V-SVZ neural cell lineage (subsetted from the merged multiomic datasets in Figures S3E-S3F) after one round of batch-correction, clustered on the basis of the transcriptional data. Distinct clusters are colored and numbered and cell types are annotated and outlined based upon marker gene expression. **(H)** UMAP of the same P2 V-SVZ cell lineage multiomic dataset shown in (G), clustered on the basis of the snATAC-seq chromatin accessibility data. Colors and numbers represent epigenetically-distinct cell clusters, and cell type annotations (indicated by the hatched lines) are based upon marker gene chromatin accessibility analysis. **(I)** UMAP showing the P6/7 V-SVZ neural cell lineage (subsetted from the merged multiomic datasets in Figures S3G-S3H) after one iteration of Harmony, clustered on the basis of the transcriptional data. Distinct clusters are colored and numbered and cell types are annotated and outlined based upon marker gene expression. **(J)** UMAP of the same P6/7 V-SVZ cell lineage multiomic dataset shown in (I), clustered on the basis of the snATAC-seq chromatin accessibility data. Colors and numbers represent epigenetically-distinct cell clusters, and cell type annotations (indicated by hatched lines) are based upon marker gene chromatin accessibility analysis. **Abbreviations:** Excit., excitatory. IPs, intermediate progenitors. NSCs, neural stem cells. OLs, oligodendrocytes. OPCs, oligodendrocyte precursor cells. Prolif., proliferative. RPs, radial precursors. TAPs, transit-amplifying precursors.

We used the same approach for the E17 cortex and obtained 4,752 paired transcriptomes and epigenomes. RNA-based clustering and marker gene analysis identified all of the expected cell types including transition precursors coexpressing RP and NSC markers (RP/NSCs), IPs and a small cluster of cells expressing markers for TAPs and neuroblasts (Figure S3C). We then subsetted and clustered only the V-SVZ lineage neural cells on the basis of the transcriptomic data (Figures 3D and S3D). There was good agreement between this analysis and the E17/18 scRNA-seq analysis (that shown in Figures 2A-2C and S2D). In particular, in both datasets some IPs and RP/NSCs were predicted to be proliferative, and both TAPs and neuroblasts were present (Figures 3D and S3D). Moreover, the E17 RP/NSCs from the multiomic analysis were transcriptionally similar to the E17/18 transition precursors defined in the scRNA-seq analysis, as shown by including them in the scRNA-seq-based single-cell correlation analysis (Figure 3E). We then reclustered the E17 V-SVZ lineage cells on the basis of chromatin accessibility rather than the transcriptional data (Figure 3F). As with the E14 multiomic dataset, the different cell types were readily defined by marker gene accessibility, but the epigenetic clustering did not distinguish proliferative from non-proliferative cells.

Finally, we acquired single-cell multiomic V-SVZ datasets at P2 and P6/7, dissecting the dorsal and immediately lateral V-SVZ. At P2, we performed two independent runs, transcriptionally-identifying all of the predicted cell types in both runs (Figures 3E-3F). At P6/7, we acquired one new P7 V-SVZ dataset (Figure S3G) and reanalyzed our previously-published P6 V-SVZ dataset (Dennis et al.,^25^ GEO: GSE255405) (Figure S3H). At both timepoints, we merged the two independent datasets, and subsetted out and reclustered V-SVZ lineage neural cells on the basis of the transcriptional data (Figures 3G-3I). Analysis of the datasets of origin (Figures S3I-S3J) showed that transcriptomes from the different datasets were well-integrated after one iteration of Harmony batch correction. At P2, as seen in the scRNA-seq analysis (Figures 1H-1I), there were many TAPs, neuroblasts, and OPCs, but not early proliferative IPs, indicating that cortical precursors were generating postnatal but not embryonic progeny (Figure 3G). Notably, none of the P2 NSCs were predicted to be proliferative (Figure S3K) and they expressed NSC but not RP marker genes, consistent with the conclusion that by P2 they had acquired an NSC state. Reclustering of the P2 V-SVZ cells on the basis of chromatin accessibility rather than transcriptional data (Figure 3H) defined the same cell types, but proliferative versus non-proliferative cells were not distinguished (Figure 3H). Similar results were obtained at P6/7 (Figures 3I-3J and S3L), although at this later timepoint there were relatively more oligodendrocytes and relatively fewer late IPs/newborn neurons.

### The transition from cortical RPs to NSCs involves a gradual epigenetic switch and an epigenetically-distinct population of E17 transition precursors

We used these single-cell multiomic datasets to ask about epigenetic changes during the RP to NSC transition. Initially, we merged the E14 to P6/7 cortical precursors as defined by the snRNA-seq data and clustered the merged dataset on the basis of the transcriptional data after performing one iteration of Harmony. UMAP visualization (Figures 4A and S4A) confirmed the scRNA-seq based conclusions. In particular, while the P2/6/7 NSCs all co-clustered transcriptionally, they were distinct from the embryonic precursors. Moreover, the E14 and E17 precursors largely segregated from each other, although some proliferating E17 precursors co-clustered with the proliferating E14 RPs (cluster 2, Figure 4A).

**Figure 4.**
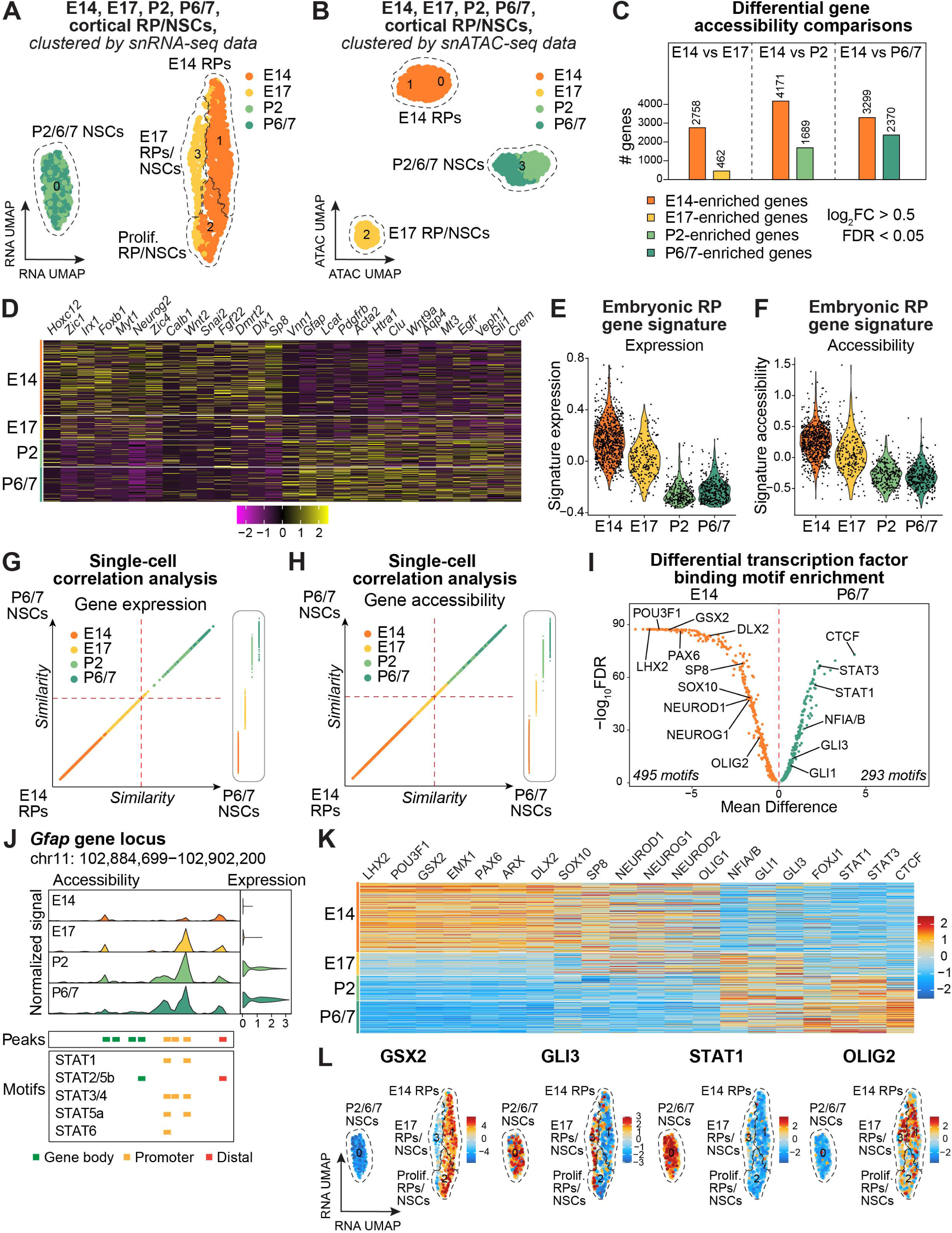
Single nucleus multiomics demonstrates that cortical precursors undergo a gradual shift in gene and transcription factor binding motif accessibility between E14 and P2. **(Also see Figure S4 and Tables S1-S3). (A)** UMAP of E14, E17, P2, and P6/7 cortical precursors from the multiomic datasets (shown in Figures 3A, 3D, 3G, and 3I) after one iteration of Harmony, clustered on the basis of the transcriptional data, annotated for cell types, and colored as per their timepoints of origin, shown in the adjacent key. Transcriptionally-distinct clusters are numbered and outlined by hatched lines (also see Figure S4A). **(B)** UMAP of the merged E14 to P6/7 cortical precursor multiomics dataset shown in (A), reclustered on the basis of the paired snATAC-seq epigenomic data and annotated for cell types. Timepoints of origin are color-coded as per the adjacent key and epigenetically-distinct clusters are numbered and outlined by hatched lines (also see Figure 4B). **(C)** Differential gene accessibility comparisons between E14 RPs versus E17 to P6/7 RP/NSCs showing the number of significantly enriched genes per comparison (FDR < 0.05 and log2FC > 0.5; Table S1). Each bar plot is color-coded based on the represented timepoints as per the adjacent key and the numbers above each bar denote the number of significantly enriched genes within each group. **(D)** Single-cell heatmap showing scaled accessibility of selected genes that were significantly enriched in a differential accessibility comparison between E14 RPs and P6/7 NSCs (Table S1). Every row represents accessibility in an individual cell, and levels are color-coded as per the adjacent key. **(E, F)** Violin plot showing mRNA expression (E) and chromatin accessibility (F) of a 15 gene embryonic cortical RP signature in E14, E17, P2, and P6/7 cortical precursors from the dataset in (A), separated by timepoint. Each dot represents gene signature levels in an individual cell. The embryonic RP signature was defined in a differential gene expression comparison between E14 RPs and P6/7 NSCs (Table S2). **(G, H)** The transcriptomes (G) and epigenomes (H) of individual RPs and NSCs from the merged E14 to P6/7 dataset shown in (A and B) were Pearson correlated against the averaged transcriptomes (G) and epigenomes (H) of E14 RPs and P6/7 NSCs. Individual cells were then plotted based on similarity to E14 RPs versus P6/7 NSCs along both x- and y-axes. Cells are colored based on timepoint as per the adjacent key. The insets on the right show the location of transcriptomes/epigenomes from each individual timepoint along the y-axis. **(I)** Transcription factor binding motif accessibility in the individual cortical precursor epigenomes shown in (B) was determined using chromVAR to determine per-cell binding motif accessibility. The transcription factor binding motifs that significantly differed were identified in a differential accessibility comparison between E14 RPs and P6/7 NSCs (Table S3) and are visualized using a volcano plot. The x-axis and y-axis indicate the log FDR (-log_2_FDR) and the mean difference in motif accessibility between the two groups, respectively. Orange and green dots denote motifs that are significantly differentially accessible in E14 RPs versus P6/7 NSCs (FDR < 0.05). **(J)** Genome browser-style plot highlighting the *Gfap* gene body and surrounding genomic regions, analyzed in the E14 to P6/7 cortical precursors shown in (B). Shown are the normalized ATAC signal within the gene locus at different timepoints (top left), violin plots showing normalized gene expression at each timepoint (top right), identified peaks within the region (middle), and peaks containing various STAT motifs (bottom). Peaks are colored based on their type according to the key within the panel. **(K)** Single-cell heatmap showing scaled enrichment scores of selected motifs identified in a differential comparison between the epigenomes of E14 RPs and P6/7 NSCs as shown in (B) (Table S3). Every column represents accessibility in an individual cell, and enrichment levels are color-coded as per the adjacent key. **(L)** UMAP of merged E14 to P6/7 cortical precursors clustered on the transcriptional data as in (A) showing accessibility enrichment z-scores for selected transcription factor binding motifs identified as being differentially accessible in a comparison of E14 versus P6/7 cortical precursors epigenomes (those shown in panel B) (Table S3), color-coded as per the adjacent keys. The motifs used in the analysis are shown in Figure S4D. **Abbreviations:** NSCs, neural stem cells. Prolif., proliferative. RPs, radial precursors.

We then reclustered the same E14-P7 dataset on the basis of chromatin accessibility. This analysis (Figures 4B and S4B) showed that as seen in the transcriptional analysis all postnatal NSCs co-clustered, the E14 and E17 precursors were completely distinct from each other at the chromatin level. We first characterized these epigenetic differences by examining global genomic accessibility, defining differentially accessible genes as having a log fold-change of greater than 0.5 (log_2_FC > 0.5) and a false discovery rate (FDR) of less than 0.05 (FDR < 0.05) (Table S1). This analysis showed that, relative to E17 transition precursors, E14 RPs were significantly more accessible at six-fold more genes (Figure 4C). However, this global difference in gene accessibility was largely reversed postnatally so that the E14 RPs were only preferentially accessible at 1.4-fold more genes than the P6/7 NSCs.

These findings suggest a model where RP gene-associated chromatin becomes less accessible between E14 and E17, coincident with an increase in accessibility for genes important for postnatal NSCs. To test this idea, we examined the individual genes involved in these changes (Table S1). This analysis demonstrated that E14 RP chromatin was preferentially open for many transcription factor genes, with the most enriched including *Hoxc12*, *Zic1*, *Irx1*, *Foxb1*, *Lhx1*, *Myt1*, *Neurog2*, *Zic4*, *Snai2*, *Hoxb2*, and *Tlx2* (Table S1). Notably, amongst these were transcription factors associated with both excitatory (*Neurog2*, *Neurod1/2/6*, *Eomes*) and inhibitory (*Dlx1/2/5*, *Sp8/9*, *Nkx2.1*) neurogenesis, suggesting that E14 cortical RPs are epigenetically primed to make inhibitory neurons even when they only generate excitatory neurons. By contrast, P6/7 NSCs were enriched for fewer transcription factor genes, and these were largely associated with signaling pathways such as *Stat2*, *Gli1*, and *Crem*. They instead showed enhanced accessibility for genes associated with the postnatal NSC state such as *Vnn1*, *Gfap*, *Pdgfrb*, *Aqp4*, *Mt3*, and *Veph1* (Table S1).

Three further analyses of gene accessibility states over time defined a gradual RP to NSC chromatin shift, with E17 transition precursors at the midpoint. First, heatmap analysis of genes preferentially accessible in E14 RPs versus P6/7 NSCs (Figure 4D) showed that the E17 transition precursors had a hybrid accessibility state that included genes that were preferentially accessible in both E14 RPs and postnatal NSCs. By contrast, P2 NSCs displayed a chromatin accessibility state very similar to P6/7 NSCs. The second analysis involved an E14 RP gene signature. Specifically, using our scRNA-seq data, we identified mRNAs that were transcriptionally enriched in E14 RPs versus P6/7 NSCs using differential gene expression analysis (Table S2) and showed that 15 of these genes were sufficient to specifically identify the E14 RPs relative to postnatal NSCs at the transcriptional level (Figures 4E and S4C; Table S2). Analysis of the chromatin accessibility data for these 15 signature genes showed that they were also more accessible in E14 RPs than in P2/P6/P7 NSCs, and that the E17 transition precursor accessibility was midway between that of the RPs and NSCs (Figure 4F).

Third, we performed single-cell correlation analysis using the merged multiomic dataset. We initially did this based on the transcriptional data, comparing each cell’s transcriptome to the averaged gene expression for E14 cortical RPs versus P6/7 NSCs. As was seen with the scRNA-seq analysis (Figure 2F), there was a continuous transition between E14 and P6/7, with the E17 transition precursors positioned midway (Figure 4G). We then did a similar analysis using the chromatin accessibility data, comparing individual epigenomes against the averaged gene accessibility scores of E14 cortical RPs and P6/7 cortical NSCs (Figure 4H). This analysis identified a gradual transition in chromatin accessibility from E14 to P6/7, with the E17 precursors located between the E14 RPs and P2 NSCs.

### The perinatal RP to NSC transition is reflected in the accessibility of regulatory transcription factor binding motifs

We took further advantage of the chromatin accessibility data to ask how the gene regulatory landscape changed during the RP to NSC transition. We did this by calculating openness for transcription factor binding motifs within each nucleus using chromVAR.^34^ A comparison of E14 RPs with P6/7 NSCs showed that the RPs were preferentially open for 495 transcription factor motifs and the NSCs for 293 (Table S3). When we examined these motifs in detail, we found that the E14 RP chromatin was preferentially open for transcription factor motifs involved in excitatory neurogenesis, including binding sites for LHX2, PAX6, NEUROD1/2, NEUROG1, POU3F1/2, and ARX^35–39^ (Figure 4I; Table S3). Unexpectedly, however, they were also highly accessible for binding motifs associated with (i) inhibitory neurogenesis such as DLX1/2 and SP8,^40,41^ (ii) oligodendrogenesis such as OLIG1/2, SOX9/10, and TCF4,^42–45^ and (iii) ventral forebrain identity such as GSX2 and NKX2.1^39,46^ (Figure 4I; Table S3). This is consistent with the data showing that the chromatin around many of the same transcription factor genes is also more accessible in E14 RPs (Figure 4D; Table S1).

By contrast, P6/7 NSC chromatin was instead more open for transcription factor binding motifs associated with downstream signaling pathways (Figure 4I; Table S3) such as for the GLIs, which are downstream of the Shh pathway. Notably, this enhanced postnatal accessibility also included motifs for the JAK-STAT signaling proteins STAT1/3 and the NFI transcription factors NFIA/B, all of which are essential for astrogenesis.^18,47,48^ Indeed, a more high-resolution analysis of STAT binding motifs within the *Gfap* gene promoter showed that they were more accessible in the postnatal NSCs than in the E14 RPs (Figure 4J), consistent with previous work showing that astrogenesis genes like *Gfap* are repressed during embryonic cortical neurogenesis.^49–51^

We asked when these differences arose by visualizing motif accessibility in individual cells within the combined cortical RP/NSC multiomic dataset (that in Figures 4A and 4B). We did this either by plotting accessibility in a heatmap (Figure 4K) or by overlaying motif accessibility on the RNA-based UMAP projection (Figures 4L and S4D-S4E). This analysis confirmed the robust differences in motif accessibility between E14 RPs and P6/7 NSCs and demonstrated that the E17 precursors were in a transition state. Some highly accessible E14 motifs were still open at E17, including those for SP8, NEUROD1/2, NEUROG1, and OLIG1, while others were already more closed, including those for GSX2, POU3F1, PAX6, and DLX2. All of the E14-enriched motifs were closed in P2 NSCs. Conversely, some P6/7 accessible motifs, such as those for GLI1, GLI3, and NFIA/B, were already becoming accessible at E17.

### Single-cell spatial transcriptomics identifies spatially-distinct embryonic cortical V-SVZ domains at E18

These data identify E17/18 as a key influx point in the transition from RPs to NSCs. To understand if this transition is spatially regulated, we performed single-cell multiplexed *in situ* gene expression analysis using the Xenium platform.^25,28,52^ We did this during the transition at E18 using a probeset targeted to 347 genes: 247 from the standard Xenium murine brain panel and an additional 100 from a custom add-on probeset that allowed better resolution of NSCs, astrocytes, and TAPs (Table S4).^25^ We analyzed coronal fresh-frozen sections at the level of the lateral ventricles, performing single-cell spatial transcriptomics as we have described.^25,28^ We defined a region of interest (ROI) that included the dorsal cortex and the top lateral corner of the lateral ventricle (Figure 5A). We analyzed one section each from 4 different mice. After initial processing of data from individual sections, we removed a small number of objects with low transcript counts (likely cellular fragments) and cellular doublets. We then merged cells from the different sections, obtaining 71,596 cellular transcriptomes.

**Figure 5.**
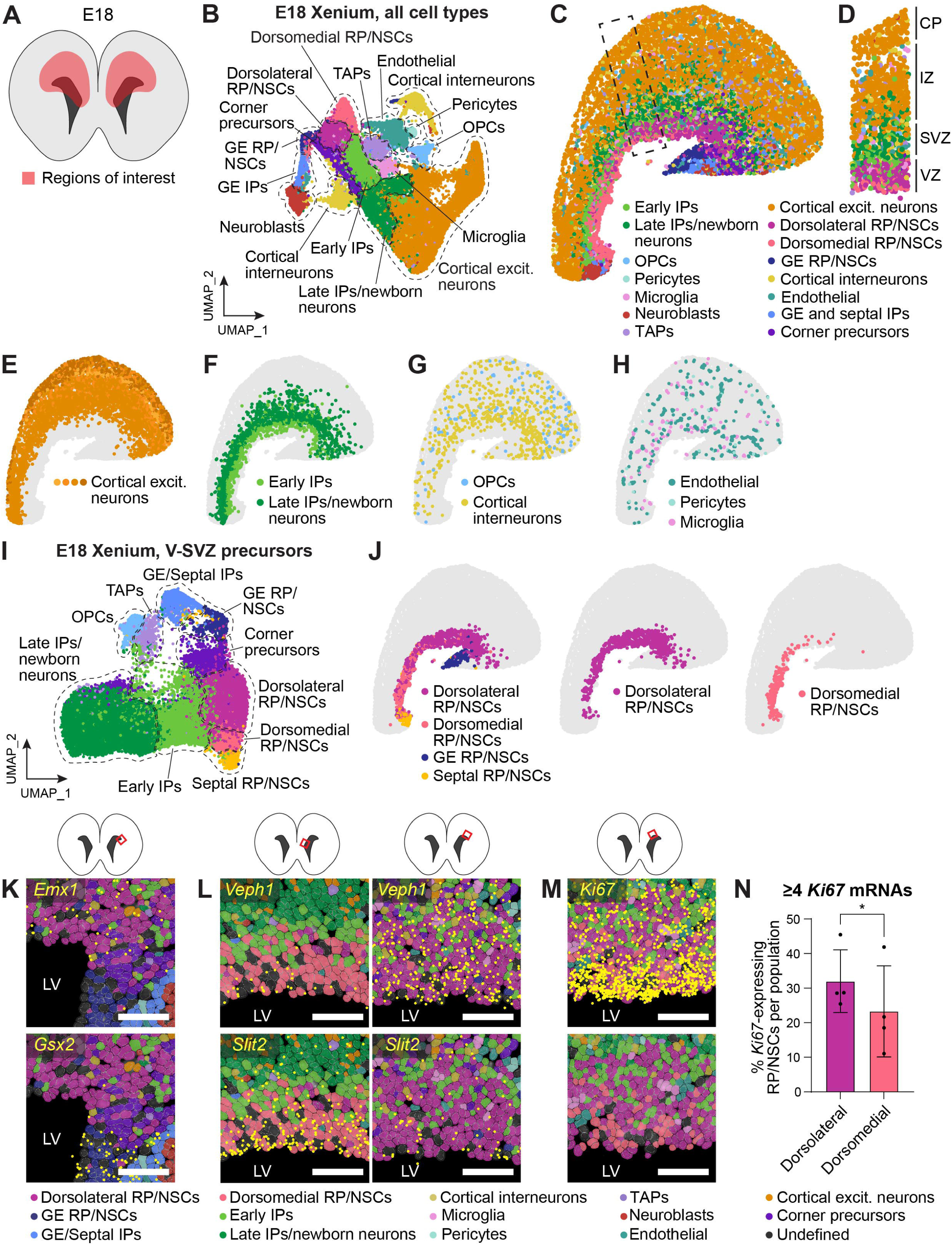
Single-cell spatial transcriptomic characterization of the E18 cortical V-SVZ identifies spatially-distinct RP/NSC populations. **(Also see Figure S5 and Tables S4-S5). (A, B)** Xenium single-cell spatial transcriptomics was performed on coronal E18 cortical sections from 4 independent animals. (A) shows a schematic of the region of interest (ROI) that was analyzed while (B) shows a UMAP of the resultant merged transcriptomes from all 4 sections/animals. Colors in (B) correspond to cell types as defined by marker gene expression. **(C)** Spatial plot showing the entire ROI from a single representative E18 hemisphere, where each cell type identified in the UMAP in (B) is color-coded as per the adjacent key. Each dot represents an individual cell. **(D)** Spatial plot of the boxed region outlined in (C) showing the different embryonic cortical layers at higher resolution. All cells are shown and are color-coded as per the adjacent key. Each dot represents an individual cell. **(E-H)** Spatial plots of the same representative E18 ROI as in (C) showing cell types plotted individually or in combination and colored as per the adjacent legends. Shown are (E) excitatory neurons, with different colors denoting spatially- and transcriptionally-distinct cortical neurons, (F) early and late IPs, (G) OPCs and cortical interneurons, and (H) microglia and vasculature-associated endothelial cells and pericytes. Each dot represents an individual cell. **(I)** UMAP of E18 V-SVZ neural cell lineage transcriptomes subsetted from the dataset shown in (B) to better define transcriptionally-distinct precursor populations. Cells are annotated and colored based upon marker gene expression and spatial locations. **(J)** Spatial plots of the same representative E18 ROI as in (C) showing RP/NSC subpopulations defined as in (I). **(K)** High-magnification Xenium Explorer images showing expression of the cortical gene *Emx1* (top panel, yellow dots) and the GE marker gene *Gsx2* (bottom panel, yellow dots) at the dorsolateral corner of the V-SVZ (schematic at the top of the panel). Also shown are the precursor cells defined in (B) color-coded as per the adjacent key. The light grey outlines denote nuclear boundaries. **(L)** High-magnification Xenium Explorer images showing expression of *Veph1* mRNA (top panels, yellow dots) and *Slit2* mRNA (bottom panels, yellow dots) in the dorsomedial V-SVZ (left panels) and dorsolateral V-SVZ (right panels). Also shown are the precursor cells defined in (B) color-coded as per the adjacent key. The light grey outlines denote nuclear boundaries, and the location of the images is shown in the schematics at the top. **(M)** High-magnification Xenium Explorer images showing expression of *Ki67* mRNA (top panel, yellow dots) in the central dorsal V-SVZ (as shown in the schematic on top). Both panels show the precursors as defined in (B), color-coded as per the adjacent key. The light grey outlines denote nuclear boundaries. **(N)** Quantification of images as in (M) for the proportion of dorsolateral versus dorsomedial RP/NSCs that express ≥ 4 *Ki67* mRNAs. Four independent sections/animals were quantified across the entire dorsal V-SVZ. n = 4, **p<0.05. **Scale bars,** 50 μm. **Abbreviations:** CC, corpus callosum. CP, cortical plate. Excit., excitatory. GE, ganglionic eminence. IPs, intermediate progenitors. IZ, intermediate zone. LV, lateral ventricle. NSCs, neural stem cells. OLs, oligodendrocytes. OPCs, oligodendrocyte precursor cells. RPs, radial precursors. SVZ, subventricular zone. VZ, ventricular zone.

UMAPs and marker gene analysis (Figures 5B and S5A-S5B) showed that the transcriptomes were well-integrated and identified all of the predicted cell types, including cortical excitatory neurons, interneurons, early IPs, late IPs/newborn neurons, OPCs, and nonneural cell types such as microglia and vasculature cells. Spatial plots demonstrated that the cortical neurons were localized to the most superficial parts of the ROI (Figures 5C-5E), as predicted. The early IPs were located in the SVZ while late IPs/newborn neurons were more superficially located in the intermediate zone (Figures 5C, 5D, and 5F), consistent with their migration into the cortical layers. Interneurons were seen throughout the ROI, albeit at a higher density in the SVZ and the intermediate zone (Figure 5G). OPCs, microglia and vasculature cells were all localized throughout the ROI (Figure 5G-5H).

This analysis also identified E18 RP/NSCs and the first cortical TAPs. To better-characterize these cell types, we subsetted and reclustered the RP/NSCs, IPs, TAPs, and late IP/newborn neurons and identified the input cells using marker genes (Figure 5I). This analysis, coupled with spatial plots, identified four spatially-distinct RP/NSC groups, all of which expressed RP/NSC genes like *Aldoc, Slc1a3, Vnn1, Tspan18*, and *Hes5* (Figure S5C). One of these included septal RP/NSCs located ventral to the medial edge of the cortical V-SVZ (Figure 5J). A second population were GE RP/NSCs that were ventral to the lateral end of the *Emx1*-positive cortical boundary and that were enriched for *Gsx2* (Figure 5J-5K). Notably, at this border between the cortex and the GE, there was also a population of cells that were highly enriched for *Ano1* and *Sema6a* (Figure S5F). We speculate that these correspond to the antihem, a region thought to be a signaling center involved in cortical specification.^53^

The two other stem cell populations were both dorsal cortical RP/NSCs that differed in their spatial location. One of these included cells that were predominantly localized more laterally, ending at the GE border (dorsolateral RP/NSCs) while the other included cells that were predominantly localized more medially, ending at the septal border (dorsomedial RP/NSCs) (Figure 5J). Regions close to the GE and septal borders were highly enriched for one of the two RP/NSC populations, but in the central region these two subpopulations were intermingled with no apparent differences in their local organization (Figures 5J, 5L, and 5M). Differential gene expression analysis showed that the dorsolateral RP/NSCs were, relative to the dorsomedial cells, enriched for *Veph1*, while the dorsomedial RP/NSCs were enriched for *Slit2*, *Gfap*, and *Rmst* (Figures 5L and S5D; Table S5). We confirmed that there was also segregation of these mRNAs within the combined E17/18 scRNA-seq dataset (Figure S5E).

We asked about proliferation of the dorsolateral versus dorsomedial RP/NSCs by characterizing expression of the proliferation gene *Ki67*. This analysis showed that *Ki67* was most robustly expressed in RP/NSCs with nuclei located immediately adjacent to the ventricle wall and that this was the case along the entire cortical V-SVZ (Figure 5M). Quantification showed that about 32% of E18 cortical RP/NSCs had robust *Ki67* expression, and that this was somewhat higher in the dorsolateral versus dorsomedial precursors (Figure 5N).

### The switch to olfactory neurogenesis first occurs in the dorsolateral cortical V-SVZ

These data identify two spatially-distinct populations of E18 RP/NSCs. We asked whether either population was associated with the onset of cortical olfactory neurogenesis. To do this, we analyzed TAPs, as identified by their enrichment for *Ascl1*, *Egfr*, *Gsx2*, and *Ki67* (Figures 5I and S5G). To ensure these were cortically-derived, we identified TAPs that also detectably expressed one or more of four genes that are expressed in the cortex but not the GE, *Emx1*, *Tfap2c, Neurod6*, or *Fezf2*. All of these mRNAs are expressed in cortical but not GE TAPs at low levels (Figure S6A; data not shown). Spatial plots demonstrated that TAPs expressing cortical marker genes comprised the majority of total TAPs in the ROI at this timepoint (Figures 6A and S6B).

**Figure 6.**
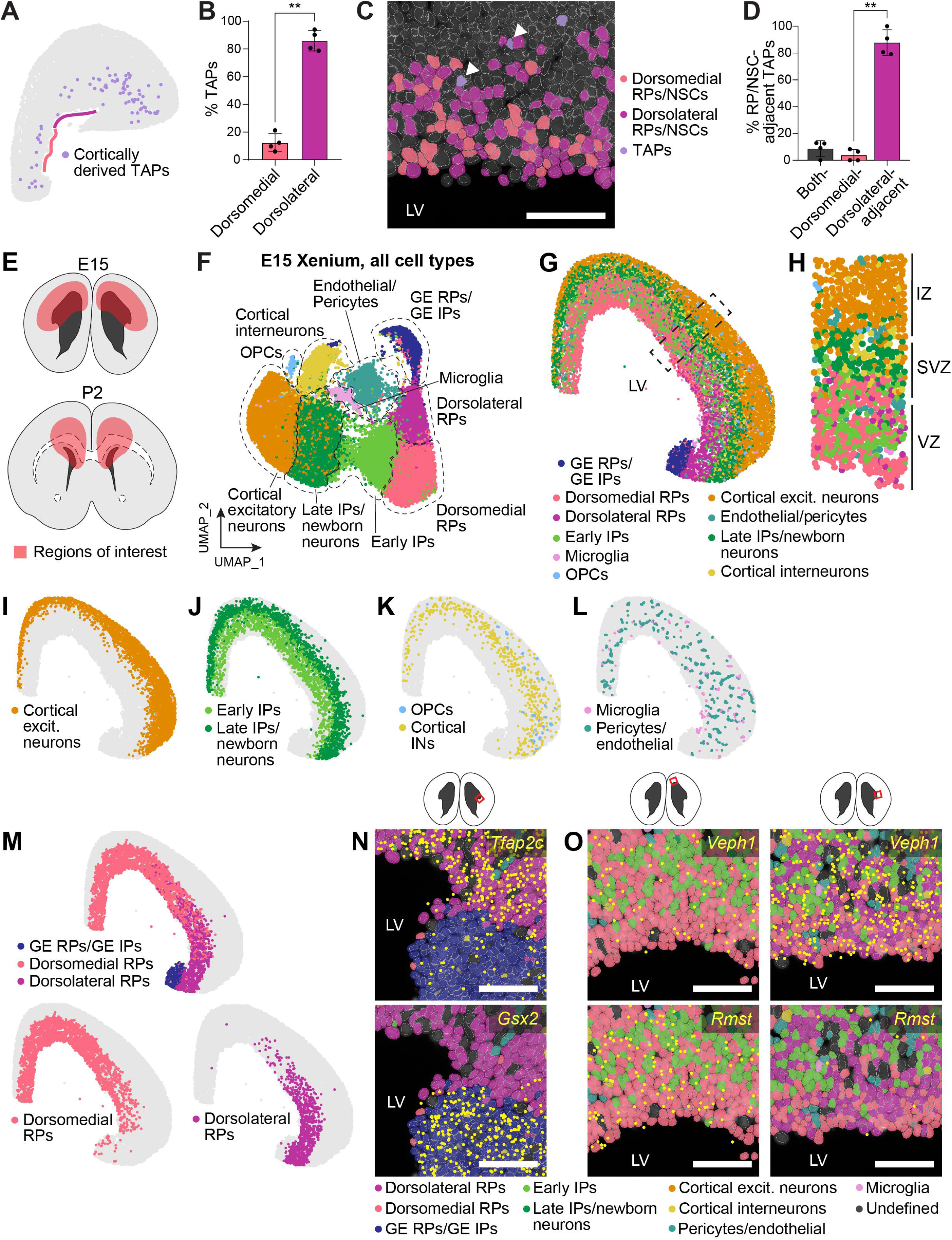
Single-cell spatial transcriptomics show that the first cortical TAPs arise dorsolaterally at E17/18, and that dorsolateral and dorsomedial RPs are also present at E15. **(Also see Figure S6). (A)** Spatial plot of cortical TAPs on the same representative E18 ROI as shown in Figure 5C. The colored lines along the ventricle border denote the dorsolateral versus dorsomedial V-SVZ regions (purple and pink, respectively). **(B)** Quantification of the proportion of cortical TAPs that are located dorsolaterally versus dorsomedially (as defined in A). n = 4, **p < 0.01. **(C, D)** Quantification of the proportion of cortical TAPs that are immediately adjacent to dorsolateral versus dorsomedial RP/NSCs across the entire dorsal V-SVZ. (C) shows a representative high-magnification Xenium Explorer image of the central cortical V-SVZ illustrating the intermingling between dorsolateral and dorsomedial RP/NSCs and the close association between the former and cortical TAPs (arrowheads). (D) shows quantification of images such as in (C). n = 4 independent animals/sections, **p < 0.01. **(E)** Schematic diagram showing ROIs analyzed for the single-cell spatial transcriptomics at E15 (top) and P2 (bottom). At both timepoints, a region encompassing the dorsal V-SVZ, part of the cortex, and some of the ventral V-SVZ was analyzed. **(F)** Xenium single-cell spatial transcriptomics was performed on 3 independent E15 coronal cortical sections (each from a different litter) as shown in (E) and the resultant transcriptomes were merged and run through the pipeline. The UMAP shows the merged transcriptomes, annotated for cell types as defined by marker gene expression. Colors correspond to annotated cell types. **(G)** Spatial plot showing the entire ROI from a single representative E15 hemisphere, where the cell types identified in (F) are color-coded as per the adjacent key. Each dot represents an individual cell. **(H)** Spatial plot of the boxed region outlined in (G) showing the different embryonic cortical layers at higher resolution. All cells are shown and are color-coded as per the adjacent key. Each dot represents an individual cell. **(I-M)** Spatial plots of the same representative E15 ROI as in (G) showing cell types plotted individually or in combination, and colored as per the adjacent legends. Shown are (I) cortical excitatory neurons, (J) early IPs and late IPs/newborn neurons, (K) OPCs and cortical interneurons, (L) microglia, vasculature-associated endothelial cells, and pericytes, and (M) transcriptionally-distinct RP populations. Each dot represents an individual cell. **(N)** High-magnification Xenium Explorer images showing expression of the cortical gene *Tfap2c* (top panel, yellow dots) and the GE marker gene *Gsx2* (bottom panel, yellow dots) at the boundary between the dorsal and ventral V-SVZ at the dorsolateral corner (schematic at the top of the panel). Also shown are the precursor cells in that region defined as in (F) and color-coded as per the adjacent key. The light grey outlines denote nuclear boundaries. **(O)** High-magnification Xenium Explorer images showing expression of *Veph1* mRNA (top panels, yellow dots) and *Rmst* mRNA (bottom panels, yellow dots) in cells of the dorsomedial V-SVZ (left panels) and dorsolateral V-SVZ (right panels) (as in the schematics on top of the panels). Also shown are the precursor cells in those regions color-coded as defined in (F) and color-coded as per the adjacent key. The light grey outlines denote nuclear boundaries. **Scale bars,** 50 μm. **Abbreviations:** CC, corpus callosum. Excit., excitatory. GE, ganglionic eminence. INs, interneurons. IPs, intermediate progenitors. IZ, intermediate zone. LV, lateral ventricle. NSCs, neural stem cells. OLs, oligodendrocytes. OPCs, oligodendrocyte precursor cells. RPs, radial precursors. SVZ, subventricular zone. TAPs, transit-amplifying precursors. VZ, ventricular zone.

Quantification showed that, of the total dorsal cortical TAPs, over 80% were located dorsolaterally (Figures 6A and 6B). Moreover, a higher resolution analysis within the cortical V-SVZ (Figure 6C) showed that cortical TAPs were very frequently associated with a dorsolateral RP/NSC, and only occasionally with a dorsomedial RP/NSC (Figure 6D), and that this was true throughout the extent of the dorsal V-SVZ. Thus, the first cortical TAPs are predominantly dorsolaterally-localized and are likely generated by dorsolateral RP/NSCs.

### Single-cell spatial transcriptomics identify two spatially-distinct E15 cortical RP populations

These findings indicate that the first cortical TAPs are generated in the dorsolateral V-SVZ. We therefore asked if we could distinguish the spatial location of the E18 RP-like precursors, transition precursors, and NSCs, and determine how this might relate to the onset of postnatal cell genesis. To do this, we compared the E18 RP/NSCs in the Xenium dataset to E15 RPs and to P2 NSCs that were also defined by single-cell spatial transcriptomics. We performed Xenium using the same probeset on one section from each of 3 different E15 or P2 mouse forebrains. We analyzed ROIs similar to those characterized at E18 (Figure 6E) and then merged all individual section datasets from a given timepoint.

Within the merged E15 dataset, marker gene analysis identified all of the predicted cell types (Figures 6F and S6C) including RPs, IPs, excitatory neurons, cortical interneurons, a small number of OPCs, and nonneural microglia and vasculature cells. Spatial plots showed that these cell types were appropriately localized (Figures 6G and 6H). Excitatory neurons were located in the cortical plate and intermediate zone, late IP/newborn neurons in the intermediate zone and early IPs in the SVZ (Figures 6G-6J). Cortical interneurons were located throughout the ROI, although they were enriched in the SVZ and intermediate zone, and OPCs were localized more laterally (Figures 6G, 6H, and 6K), consistent with their recent migration into the cortex from the GE. Vasculature cells and microglia were distributed throughout the ROI (Figure 6L).

Importantly for our analyses, there were three groups of RPs, all expressing the RP/NSC mRNAs *Aldoc*, *Slc1a3/Glast*, *Tspan18*, and *Hes5*, as well as the proliferation mRNA *Ki67* (Figures 6F and S6C-S6D). Spatial plots demonstrated that one group included *Gsx2*-positive GE RPs located immediately ventral to the *Tfap2c*-positive cortical boundary on the lateral V-SVZ (Figure 6M and 6N). The other two groups were spatially analogous to the dorsolateral and dorsomedial RP/NSCs seen at E18 (Figure 6M). These two E15 RP populations overlapped in the central V-SVZ, but one extended laterally to the GE boundary while the other extended more medially (Figure 6M and 6O). The dorsolateral RPs were enriched for *Mt3*, *Tox*, and *Veph1* while the dorsomedial were enriched for *Slit2* and *Rmst* (Figures 6O and S6D), as was seen at E18. Moreover, also as seen at E18, the cortical cells immediately adjacent to the GE were enriched for *Ano1* and potentially comprise the antihem (Figure S6D). We confirmed segregation of the dorsolateral versus dorsomedial RP mRNAs within our E14 multiomic dataset; on both the RNA and chromatin accessibility levels, *Rmst* was enriched in a different group of RPs than were *Veph1* and *Tox* (Figure S6E and S6F). Thus, at both E15 and E18, there are two spatially-distinct dorsal cortical precursor groups.

### At P2, non-proliferative NSCs and proliferative TAPs are located along the entire dorsal V-SVZ

A similar analysis at P2 identified all of the cell types predicted by our scRNA-seq and multiomic analyses; cortical excitatory and inhibitory neurons, late IPs/newborn neurons, TAPs, olfactory neuroblasts, astrocytes, OPCs, microglia, and vasculature cells (Figure 7A). Spatial plots showed these were all appropriately localized (Figures 7B and 7C). Excitatory neurons were located in the cortical layers while late IP/newborn neurons were apparently migrating from the SVZ to the cortical layers (Figures 7B-C and S7A). Cortical interneurons were located within the cortical parenchyma, OPCs and astrocytes were located in the parenchyma and nascent corpus callosum, and newborn oligodendrocytes were enriched in the corpus callosum (Figures 7B-7C and S7B-S7D). Vasculature cells were located throughout the ROI (Figure S7E). Notably, at this age, microglia were enriched within the SVZ/nascent corpus callosum (Figure S7D), consistent with a potential role in regulating oligodendrogenesis.^28,54^

**Figure 7.**
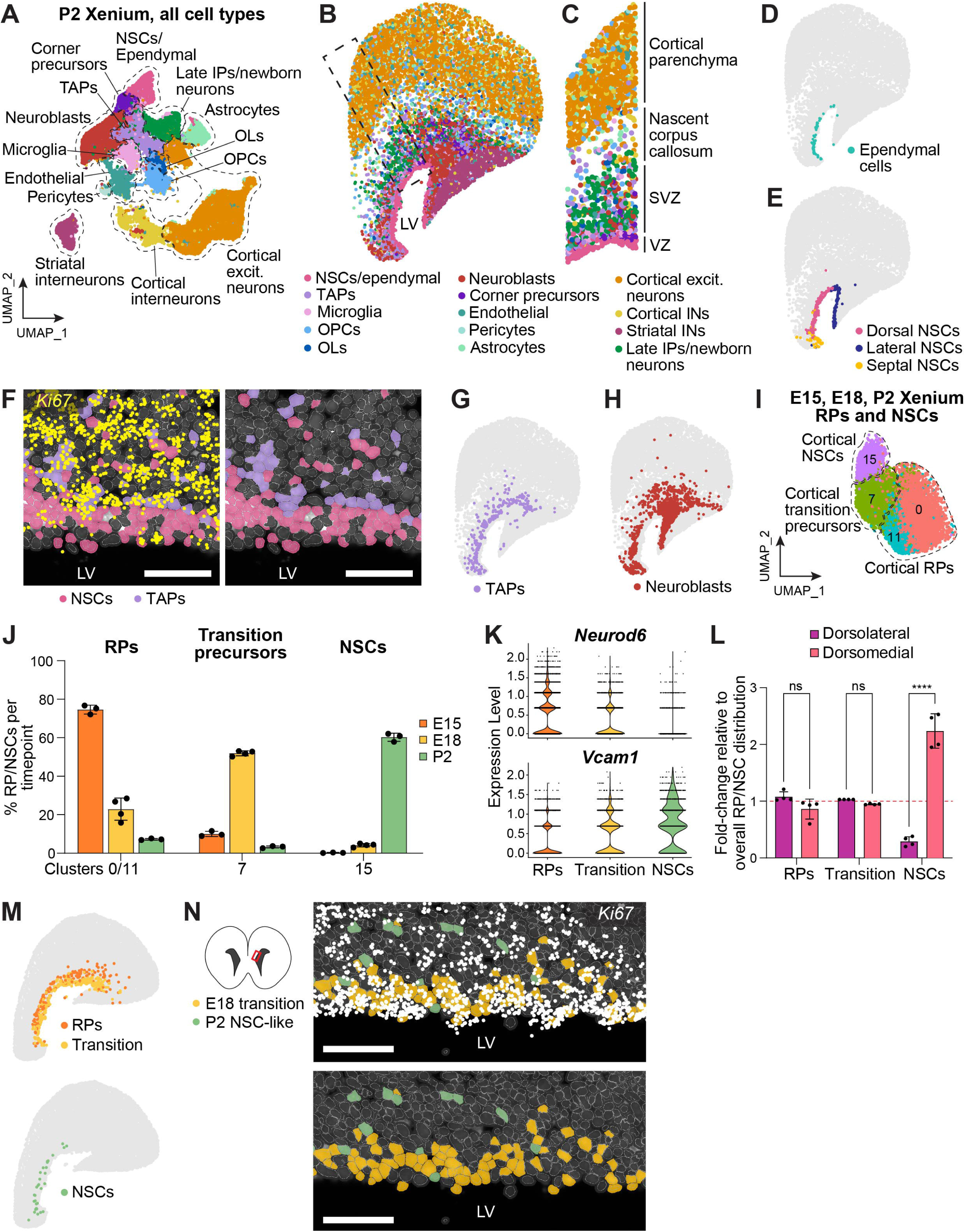
Single-cell spatial analysis identifies the location of E18 transition precursors and shows that the first cortical NSCs arise dorsomedially. **(Also see Figure S7). (A)** Xenium single-cell spatial transcriptomics was performed on 3 independent P2 coronal cortical sections (each from an independent animal) and the transcriptomes from each individual ROI (as in Figure 6E) were merged and run through the pipeline. The UMAP shows the merged transcriptomes, annotated for cell types as defined by marker gene expression. Colors correspond to annotated cell types. **(B)** Spatial plot showing the entire ROI from a single representative P2 hemisphere, where the cell types identified in (A) are color-coded as per the adjacent key. Each dot represents an individual cell. **(C)** Spatial plot of the boxed region outlined in (B) showing the different cortical layers at higher resolution. All cells are shown and are color-coded as per the adjacent key. Each dot represents an individual cell. **(D, E)** Spatial plots of the same representative P2 ROI as in (B) showing (D) ependymal cells and (E) cortical, GE/lateral, and septal NSCs colored as per the adjacent legend. Cell types are annotated as per the UMAP in Figure S7F. **(F)** High-magnification Xenium Explorer images showing expression of *Ki67* mRNA (left panel, yellow dots) in the central dorsal V-SVZ. In both panels, cell types are annotated as in Figure S7F and color-coded as per the adjacent key. The light grey outlines denote nuclear boundaries. **(G, H)** Spatial plots of the same representative P2 ROI as in (B) showing (G) TAPs and (H) neuroblasts. Cell types are annotated and color-coded as per the UMAP in (A). Each dot represents an individual cell. **(I)** UMAP as in Figure S7I showing only the clusters containing the majority of E15 cortical RPs (clusters 0 and 11), E18 cortical transition precursors (cluster 7), and P2 cortical NSCs (cluster 15). The transcriptionally-distinct clusters are colored and numbered. Within each cluster, cells relatively far outside of the cluster boundaries were removed for visual clarity. **(J)** Quantification of the proportion of cortical precursors from each timepoint that localize to the RP clusters (0 and 11), transition precursor cluster (7) or NSC-like cluster (15) in the merged dataset shown in panel I and Figure S7I. Bars are colored based on timepoint and each dot represents a different animal (E15, n = 3; E18, n = 4, P2, n = 3). Error bars denote standard deviation. **(K)** Violin plots showing the normalized expression of the RP marker gene *Neurod6* and the NSC gene *Vcam1* in the RPs (clusters 0/11), transition precursors (cluster 7), and NSCs (cluster 15) shown in (I) and Figure S7I. Each dot represents expression levels in an individual cell. **(L)** Quantitation of the relative distribution of RPs, transition precursors and NSCs in the dorsolateral versus dorsomedial domains of the E18 dorsal V-SVZ (see Figure 6A). Proportions were normalized to the distribution of all precursors in each domain. Each dot represents a different animal (n = 4), and error bars denote standard deviation. ns = nonsignificant and ****p < 0.001. **(M)** Spatial plots of the same representative P2 ROI as in (B) showing RPs, transition precursors and NSCs as defined in the UMAP in (I). Cell types are colored as per the adjacent legends. **(N)** High-magnification Xenium Explorer images of the dorsomedial cortical V-SVZ showing expression of *Ki67* mRNA (top panel, white dots) in transition precursors (yellow) and NSCs (green). **Scale bars,** 50 μm. **Abbreviations:** Excit., excitatory. INs, interneurons. IPs, intermediate progenitors. LV, lateral ventricle. NSCs, neural stem cells. OLs, oligodendrocytes. OPCs, oligodendrocyte precursor cells. RPs, radial precursors. SVZ, subventricular zone. TAPs, transit-amplifying precursors. VZ, ventricular zone.

This dataset also included NSCs. To ensure we could distinguish these from ependymal cells, which are just starting to be generated in the cortex at P2, we subsetted out and reanalyzed the NSCs, ependymal cells, TAPs, IPs, and early neuroblasts (Figure S7F). This analysis identified *Foxj1*-high ependymal cells that only partially lined the cortical VZ (Figures 7D and S7F), consistent with previous reports.^55^ There were also three groups of NSCs that expressed the RP/NSC mRNAs *Aldoc*, *Slc1a3/Glast*, *Hes5*, and *Tspan18*. One included GE/lateral wall NSCs expressing *Gsx2* but not the cortical markers *Tfap2c* or *Emx1* (Figures 7E and S7G). A second group included septal NSCs located on the medial wall (Figure 7E). The third included cortical NSCs that extended the length of the dorsal ventricle wall between the GE and septal NSCs, and that expressed *Tfap2c* and *Emx1* but not *Gsx2* (Figures 7E and S7G). We did not observe distinct lateral and medial populations along the dorsal wall at this age. Moreover, higher resolution analysis showed that almost all dorsal cortical NSCs were *Ki67*-negative (Figure 7F), consistent with the idea they have acquired an NSC state and commenced the transition to dormancy, as seen in the scRNA-seq datasets. Thus, by P2, dorsal cortical NSCs are relatively homogeneous and non-proliferative, consistent with the scRNA-seq and multiomic data.

This analysis also demonstrated that, relative to E18, the situation had evolved with regard to cell genesis. In particular, TAPs were more numerous than at E18 and were located all along the dorsal V-SVZ rather than just in the dorsolateral domain (Figure 7G). By contrast to the NSCs, these TAPs were almost all *Ki67*-positive (Figure 7F). There were also many medially-localized olfactory neuroblasts in addition to those located more dorsolaterally and in the rostral migratory stream (Figure 7H). These were presumably generated by the medially-localized P2 TAPs. Finally, we also defined a group of dorsolateral corner precursors that extended into the adjacent cortical parenchyma (Figures 7A and S7H). These corner precursors expressed both NSC genes and lower levels of neurogenic TAP genes such as *Dlx1*, *Dlx2*, and *Gad1*, and we speculate they may be early neurogenic TAPs.

### At E18, transition precursors are distributed along the entire cortical V-SVZ, whereas the first non-proliferative NSCs are located medially

We then used these E15 and P2 Xenium datasets to define the transcriptional states and spatial locations of E18 RP/NSCs. Specifically, we subsetted and merged RP/NSCs, TAPs, IPs, and late IP/newborn neurons from the E15, E18, and P2 Xenium datasets. Analysis of this merged dataset identified all of the input cell types (Figure S7I) and demonstrated that the majority of RP/NSCs were in 3 groups (Figure 7I). One of these (cluster 15, termed NSCs) included most P2 dorsal cortical NSCs and a small proportion of E18 RP/NSCs (Figure 7J). A second transcriptionally-distinct cluster (7, termed transition precursors) included most E18 RP/NSCs and small proportions of E15 and P2 precursors (Figure 7J). Finally, a third group (clusters 0 and 11, termed RPs) included most E15 RPs and about 20% of the E18 RP/NSCs (Figure 7J), consistent with the scRNA-seq analysis (Figures 2D-2F and 2H). We validated these assignments by analyzing expression of Xenium probeset genes that distinguish RPs and NSCs. Relative to the RPs, the NSCs were significantly enriched for *Vcam1*, *Vnn1*, *Aqp4*, *Thbs4*, *Slc1a3*, *Clu*, *Mdfi*, *Gfap*, and *Ttyh1*, all known NSC genes (Figures 7K and S7J). Conversely, the RPs were significantly enriched for *Neurod6*, *Eomes*, *Cdh6, Ki67*, and *Fn1*, genes associated with excitatory neurogenesis, cell adhesion, neuroepithelial function, or proliferation (Figures 7K and S7J). Notably, as seen in the other analyses, the transition precursors expressed intermediate levels of many of these genes (Figures 7K and S7J).

These spatial transcriptomic findings indicate that most E18 RP/NSCs identified in the Xenium analysis are transition precursors, some are RPs, and a small number are NSCs, in excellent agreement with the other analyses. We therefore asked about the spatial location of these different E18 RP/NSC populations, quantifying their dorsomedial versus dorsolateral distributions relative to the total RP/NSCs in those domains. This analysis showed that the transition precursors were evenly distributed along the dorsal V-SVZ, as were the less-abundant RPs (Figures 7L and 7M). By contrast, the NSCs were not evenly distributed but were instead highly enriched in the dorsomedial domain (Figures 7L and 7M). A higher resolution analysis defined two additional differences between the E18 transition precursors and the NSCs. First, in locations where both precursor types were present, the transition precursors were often located closer to the ventricle wall than were the NSCs (Figure 7N). Second, many of the transition precursors expressed *Ki67* mRNA, but the NSCs were virtually all negative for this proliferative mRNA (Figure 7N). Thus, at E18 the first non-proliferative cortical NSCs are dorsomedially-localized, while the first cortical TAP genesis occurs dorsolaterally, indicating that NSCs are not the source of cortical olfactory bulb neurons at this stage, and implying that the acquisition of an NSC state and the switch in cell genesis are separable transitions.

## Discussion

Here we have asked how the adult cortical NSC pool is established and how this relates to the development switch from making embryonic excitatory neurons to generating postnatal glial cells and interneurons. Regarding NSC establishment, we find that proliferative embryonic cortical RPs undergo a gradual transcriptional and epigenetic transition to become non-proliferative NSCs postnatally. These findings support a continuous model of cortical NSC formation in contrast to the set-aside model proposed for GE-derived NSCs. In this continuous model, as excitatory neurogenesis declines during mid- to late-embryogenesis, highly-active cortical RPs start to acquire epigenetic and transcriptional characteristics of an NSC state while at the same time losing their characteristic RP state. By E17/18, these cortical precursors are in a unique transition state where they are epigenetically open for and express both RP and NSC genes. These transition precursors are proliferative and as a population make both embryonic and postnatal progeny. By P2, the transition precursors have become non-proliferative NSCs and, while not studied here, these newly-generated NSCs will subsequently undergo a transition to full dormancy, as we have previously described.^4^

How does this RP to NSC transition relate to the shift from making embryonic to postnatal cell types? Our data indicate that these two transitions are dissociable and that cortical precursors start to make their postnatal progeny before they become NSCs. In particular, we show that, at E18, cortical TAPs are first seen dorsolaterally while the first NSCs present at this timepoint are located dorsomedially, indicating that postnatal cell types are being made by the transition precursors. Moreover, we provide evidence that embryonic cortical RPs are epigenetically primed to make both embryonic and postnatal progeny, consistent with previous work showing that embryonic cortical precursors will make glia and interneurons when their local environment is changed by transplantation, exogenous ligand exposure, or manipulation of downstream signaling pathways.^13,18,19,56–58^

Together these findings indicate that embryonic cortical precursors are multipotent, and that genesis of embryonic versus postnatal progeny does not require acquisition of an NSC state. Instead, we posit that the cell genesis switch is determined at least in part by the local environment. What then is different about the dorsolateral V-SVZ niche that might cause genesis of TAPs specifically in this domain? We propose that one key difference involves GE-derived cortical interneurons; our spatial transcriptomic data identify an increased density of these cells in the dorsolateral domain perinatally (see Figure 5G), and secretion of fractalkine and Shh by these interneurons has been implicated in the initial cortical genesis of both OPCs and olfactory neuroblasts.^12,13,19^

If transition RP/NSCs make glia and interneurons during late embryogenesis, then what happens postnatally when there are only NSCs? In adults the lineage is very clear; dormant forebrain NSCs are activated to generate TAPs and these make glia and olfactory interneurons. However, the situation is less clear in the early postnatal period when there is a requirement for high levels of cell genesis at the same time that cortical NSCs need to become fully dormant to establish the adult NSC pool. One possible explanation is that there is a lineage bifurcation that occurs around E17/18, with some transition precursors becoming NSCs that progress to full dormancy, and others becoming multipotent proliferative TAPs that can expand and support early postnatal cell genesis. Our P2 clustering and trajectory analyses support this idea (see Figure 1J), with neonatal TAPs rather than NSCs at the hub of the differentiating glial and interneuron lineages. Further support comes from a recent paper that identified a transient pool of proliferative postnatal cortical precursors that generates both interneurons and glial cells.^20^ Nonetheless, even if this is the case perinatally, ultimately there must be a switch to the adult lineage where cell genesis depends upon activation of dormant NSCs. Perhaps this switch occurs after P15 when NSCs have attained their final adult transcriptional state.^4^

How do our findings relate to the previously proposed set-aside model? This model is based upon label-retaining studies that identified a population of slowly-proliferating, set-aside embryonic RPs that generate the adult lateral V-SVZ NSC pool.^5,6^ Notably, these studies did not identify set-aside RPs within the embryonic cortex, and did not further address the origins of dorsal V-SVZ NSCs. Moreover, a later study found that adult label-retaining dorsal NSCs derive from precursors that are still proliferating perinatally,^7^ consistent with our finding that most dorsal NSCs arise between E18 and P2. Why would these two pools of V-SVZ NSCs be generated by such different mechanisms? It may be that the set-aside model makes biological sense for the GE, where cell genesis occurs embryonically and is temporally separated from the postnatal establishment of the NSC pool. By contrast, the continuous model we propose here makes more sense for the cortex where these two events occur within the same perinatal timeframe. The cortex may instead be more similar to the hippocampal dentate gyrus, where granule cell genesis and establishment of the NSC pool both occur postnatally, and where the latter has also been proposed to occur via a continuous model.^3^

One unexpected finding reported here is the presence of two spatially-segregated and transcriptionally-distinct populations of cortical precursors in the embryonic dorsal V-SVZ. At E15 and E18, we find that rather than constituting uniform populations, cortical precursors separate into dorsomedial and dorsolateral populations. The medial and lateral extremities of the dorsal V-SVZ contain predominantly one or the other population, while in the middle, the two intermingle. Notably, several genes distinguishing these two precursor groups have been previously reported as enriched medially or laterally. One example is *Sp8* mRNA, which is enriched in the dorsomedial precursors; SP8 transcription factor protein is also enriched in the medial cortex, and is thought to play a key role in cortical patterning.^59^ A second example is *Tox* mRNA, which is enriched in dorsolateral precursors; a previous study identified enrichment of TOX transcription factor protein levels in the lateral aspect of the embryonic cortical V-SVZ.^60^

What then do these two spatially-distinct stem cell populations represent? Previous work has identified medial-lateral gradients within the embryonic murine cortex involving transcription factors such as *Pax6*, *Sp8*, *Emx2*, and *Nr2f1* and these have been implicated in establishing telencephalic identity and determining neurogenic potential.^61^ However, these gradients are thought to be important earlier in embryogenesis, whereas the dorsolateral and dorsomedial precursors we document here are seen at E15 and E18. Nonetheless, perhaps these two populations are established earlier during patterning and then maintained until the precursors become NSCs and/or excitatory neurogenesis is over. Whether these transcriptional differences result in functional differences is unclear, but it is intriguing that the first TAPs likely originate from dorsolateral precursors, while the first NSCs arise dorsomedially.

In summary, our data indicate that embryonic RPs give rise to postnatal NSCs as a population via a gradual transcriptional and epigenetic transition during the developmental window surrounding birth, and this transition is dissociable from the transition to generating postnatal cell types. One final open question involves the specific mechanisms that drive the transition from proliferative cortical RPs to non-proliferative postnatal NSCs. Our data do not answer this question, but do support the idea that this process involves a close interplay between an evolving cortical environment and the precursor epigenetic state. Ultimately, answering this question will provide insights into the complex interplay of factors that dictate proper establishment of adult NSC pools, and may define strategies for promoting brain repair by allowing adult NSCs to reacquire their embryonic multipotency.

## Acknowledgements

This work was funded by CIHR (to DRK and FDM). We thank Dr. Jasmine Yang for her technical assistance.

## Author contributions

BSW conceptualized, performed and analyzed experiments and co-wrote the manuscript. KK and NT collected P2 *Emx1-Eyfp* scRNA-seq and P2 V-SVZ multiomic data. DJD collected E18 cortical scRNA-seq data. BSW performed all other experiments and analyses in the manuscript. FDM and DRK conceptualized experiments, analyzed data, and co-wrote the manuscript.

## Materials and Methods

### Resource Availability

#### Lead contact

Further information and requests for resources and reagents should be directed to and will be fulfilled by the lead contact, Freda Miller.

#### Data availability

All raw scRNA-seq, multiomic, and Xenium datasets will be made available on GEO upon publication.

### Experimental Model and Subject Details

#### Mice

All animal use was approved by the Animal Care Committees of either the Hospital for Sick Children or the University of British Columbia in accordance with the Canadian Council of Animal Care policies. Mice were fed rodent chow and had free access to water in a 12-hour dark-light cycle room. All mice were well maintained in a healthy state and no mouse displaying any signs of a health or behavioral abnormality was used in the study.

Wild-type CD1 and C57BL/6 mice were obtained from Charles River Laboratories. CD1 mice were used for the E18 cortex scRNA-seq dataset and the E15 and E18 Xenium experiments. C57BL/6 mice were used to generate all multiomic datasets (E14, E17, P2, and P6/7), as well as the P2 Xenium data. Brains for the E15 Xenium data were collected from pregnant mice that had been intraperitoneally administered 200 μL of 0.1% BSA (Jackson) in PBS 48 hours prior to collection.

*Emx1Cre* (B6.129S2-Emx1^tm1(cre)Krj^/J, RRID: IMSR_JAX:005628)^23^ and *R26Eyfpfl/fl* (B6.129X1-61 Gt(ROSA)26Sor^tm1(EYFP)Cos^/J, RRID: IMSR_JAX: 006148)^62^ transgenic mouse lines were obtained from Jackson Laboratories. All transgenic mice were bred and genotyped as recommended by Jackson Laboratories. For lineage-tracing experiments, *Emx1Cre* mice were crossed with *R26Eyfpfl/fl* mice, and resultant *Emx1Cre;R26Eyfpfl/fl* mice (also referred to as *Emx1-Eyfp* mice) were used. *Emx1Cre;R26R-Eyfp* mice were used for the P2 *Emx1-Eyfp* scRNA-seq dataset, as well as the P2 *Emx1-Eyfp* immunohistological analysis.

For all studies, mice of either sex were used. The age of the mice used in the study ranged from embryonic day 14 (E14) to postnatal day 7 (P7). The specific ages of each animal used for specific experiments is documented within the results, in the relevant section.

## Method Details

### Single-cell isolation & 10x Genomics

For single-cell isolation of the embryonic (E14, E17, E18) cortex, the meninges were removed and tissue immediately dorsal to the lateral ventricles—including the cortical layers and the V-SVZ—was collected. For the E18 cortex scRNA-seq dataset, thick coronal sections rostral to the hippocampus were created and the tissue dorsal to the lateral ventricles was dissected out. For single-cell isolation of the postnatal (P1, P2, P6, P7) V-SVZ, the cortex was removed to expose the lateral ventricles, and the dorsolateral V-SVZ was microdissected out and collected. During dissections, collected tissue was placed in either DMEM (Gibco) or Hibernate A without calcium (BrainBits) supplemented with 2% B27 (Life Technologies) and 500 µM L-glutamine (Cedarlane) and left on ice until subsequent processing steps.

For some datasets (E18 cortex scRNA-seq, E17 cortex multiome, P2 V-SVZ n2 multiome, P7 V-SVZ multiome), tissue collection was immediately followed by an enzymatic digestion step. Samples were incubated in 20 units/mL Papain with 1 mM L-cysteine, 0.5 mM EDTA, and 0.005% DNase I (Worthington) dissolved in Hibernate A medium without calcium (BrainBits) and DRAQ5 (Abcam) diluted 1:3000 for 30 minutes at 37°C to simultaneously enzymatically digest the tissue and label nuclei. Following enzyme treatment, cells were spun down at 300 g for 5 minutes and resuspended in media, either DMEM (Gibco) or Hibernate A medium without calcium, supplemented with 2% B27 (Life Technologies) and 500 µM L-glutamine (Cedarlane).

Following enzymatic digestion or immediately following collection, tissue was triturated using blunt needles with sequentially decreasing diameters (McMaster-Carr). Cells were then spun down at 300 g for 5 minutes and resuspended in 0.25% BSA (Jackson) in HBSS, after which the cell suspension was filtered through a 40 µm cell strainer and incubated in 1:1000 propidium iodide (PI) (Abcam) to label dead/damaged cells. Samples were then FACS sorted using a MoFlo Cell Sorter (Beckman Coulter) or Aria Fusion A (Becton Dickinson) to enrich for viable (PI-negative, DRAQ5-positive) cells, which were collected in media, either DMEM or Hibernate A without calcium, supplemented with 2% B27 and 500 µM L-glutamine.

Single-cell capture and cDNA library preparation were then performed at the Princess Margaret Genomics Centre (Toronto, ON) using the relevant 10x Genomics Chromium system protocol. Libraries were then sequenced using Illumina HiSeq 2500 (E18 cortex scRNA-seq), Illumina NovaSeq 6000 (P2 Emx1 scRNA-seq, P2 V-SVZ n1 multiome, and P7 V-SVZ multiome), and Illumina NovaSeq X (E14 cortex multiome, E17 cortex multiome, P2 V-SVZ n2 multiome) systems.

For wild-type (CD1 or C56BL/7) scRNA-seq samples, FASTQ reads were aligned to the wild-type mouse genome (mm10). For *Emx1-Eyfp* lineage-traced scRNA-seq samples, FASTQ reads were aligned to a custom reference genome containing the *Eyfp* open reading frame concatenated to the Ensembl mm10 reference genome. All scRNA-seq datasets were processed using 10x Genomics Cell Ranger v4 software (cellranger count function) with settings as recommended by the manufacturer.

For multiomic samples, FASTQ reads were processed and aligned to the wild-type mouse genome (mm10) using 10x Genomics Cell Ranger ARC v2 software (cellranger-arc count function) with settings as recommended by the manufacturer.

### scRNA-seq data analysis

10x Genomics scRNA-seq data was analyzed using a previously described computational pipeline.^4,24–29,63,64^ In brief, cells with low unique molecular identifier (UMI) counts, red blood cells, cells with relatively high mitochondrial DNA content, and putative doublets were removed. Genes detected in fewer than three cells were removed and cell transcriptomes were normalized using the scran R package to account for read depth and library size.^65^ Cyclone was used to predict cell cycle phase.^32^ The Seurat package (v5.2.0)^30^ was then used to process the normalized expression matrices. Following this, principal component analysis (PCA) was performed using the top 2,000 highly variable genes. The 30 top principal components were used (i) to project the dataset into two-dimensional space as Uniform Manifold Approximation and Projection (UMAP) embeddings using the Seurat RunUMAP, and (ii) to iteratively carry out SNN-Cliq-inspired clustering using the Seurat FindClusters function, at resolutions ranging from 0.4 to 2.4. Each dataset was analyzed by choosing the most conservative resolution at which cell populations of known identity separated into distinct clusters; resolution was only increased to interrogate cell heterogeneity. For further information on the resolutions used within this paper, see *scRNA-seq resolutions used for data visualization* below.

Cell types were identified based on expression of canonical markers. We used the following gene expression definitions: embryonic radial precursors (RPs), postnatal neural stem cells (NSCs), and astrocytes expressed *Aldoc*, *Dbi*, *Fabp7*, *Hes1*, *Hes5*, and *Slc1a3*; postnatal NSCs expressed *Mdfi*, *Shroom3*, *Tfap2c*, *Tspan18*, *Veph1*, and *Vnn1*; astrocytes expressed *Agt*, *Aqp4*, *Hbegf*, and *Htra1*; all intermediate progenitors (IPs) expressed *Eomes*; early IPs expressed *Insm1* and *Neurog1* in addition to *Eomes* and were negative for late IP markers; late IPs expressed *Neurod1* and *Tbr1* in addition to *Eomes*; cortical excitatory neurons expressed combinations of *Cux2*, *Fezf2*, *Ldb2*, *Pcp4*, *Satb2*, *Slc17a6*, *Tbr1*, and *Unc5d*; transit-amplifying precursors (TAPs) expressed *Ascl1*, *Egfr*, and *Gsx2*; olfactory bulb neuroblasts expressed *Dlx1*, *Dlx2*, *Gad1*, *Gad2*, *Sp8*, and *Sp9*; oligodendrocyte lineage cells expressed *Olig2* and *Sox10*; oligodendrocyte precursor cells (OPCs) expressed *Cspg4* and *Pdgfra*; homeostatic OPCs (hOPCs) expressed *Npas1*, *Pcp4*, *Pnlip*, *Ptprn*, and *Hes1* in addition to OPC markers; active OPCs (actOPCs) expressed *Ascl1*, *Dll3*, *Slc38a1*, and *Nfix* in addition to OPC markers; immature oligodendrocytes expressed *Enpp6*, *Fyn*, and *Gpr17*; mature oligodendrocytes expressed *Mag*, *Mbp*, *Mog*, *Opalin*, and *Trf*; striatal interneurons expressed *Gad1*, *Ppp1r1b*, *Bcl11b*, and *Calb1*; cortical interneurons expressed *Sst*, *Pvalb*, or *Vip* in addition to *Gad1*; Cajal-Retzius cells expressed *Cacna2d2*, *Ebf3*, *Tmem163*, and *Trp73*; pericytes expressed *Cspg4*, *Pdgfrb*, *Des*, and *Rgs5*; endothelial cells expressed *Pecam1*, *Esam*, and *Plvap*; ependymal cells expressed *Acta2*, *Ccdc153*, *Foxj1*, and *Rarres*; microglia expressed *Cd68*, *Cx3cr1*, and *Aif1*; and proliferating cells expressed *Ki67* and *Top2a*. Cells expressing markers like *Gad1/2*, *Dlx1/2*, and *Sp9* in the absence of *Sp8* and cortical or striatal interneuron markers were broadly categorized as interneurons.

UMAPs and gene expression overlays, generated using the DimPlot and FeaturePlot functions, were used to investigate gene expression and interpret cell clustering patterns. Heatmaps and violin plots were generated using the DoHeatmap and VlnPlot functions, respectively.

To merge or subset datasets, barcodes corresponding to the cells or cell types of interest were used to extract relevant gene expression information from filtered gene expression matrices. For the merging of datasets, the gene expression information from each respective dataset was then combined and the cells were then run through the pipeline as described above.

### Batch correction of scRNA-seq data

Batch correction of scRNA-seq data was performed using the Harmony package in R (v1.2.0).^31^ In brief, the RunHarmony function was used to reduce batch effects that manifested between cells originating from different datasets within a Seurat object. All scRNA-seq datasets were assigned to the following batches, which largely corresponded to when and by whom each dataset was generated, and run through Harmony accordingly. Batch 1 consisted of all datasets from Borrett et al:^4^ the E14 *Emx1-Eyfp* datasets, the E17 *Emx1-Eyfp* dataset, the P2 *Emx1-Eyfp* (n1) dataset, the P6 *Emx1-Eyfp* dataset, and the P7 *Emx1-Eyfp* dataset. Batch 2 consisted of all datasets from Dennis et al.,^25^ as well as any additional cortical datasets: the E18 cortex dataset, the P2 cortex dataset, and the P7 cortex dataset. Finally, Batch 3 consisted of the P2 *Emx1* (n2) dataset. The minimum number of Harmony iterations (either 1 or 2) required for the datasets to intermingle as expected were performed; the number of Harmony iterations used per dataset is described where relevant in the results section of this paper. After using RunHarmony, new UMAP embeddings were generated using the RunUMAP function, and the data was reclustered using the FindNeighbors and FindClusters functions at a range of resolutions.

### Resolutions used for scRNA-seq data annotation and visualization

The following resolutions were used to visualize scRNA-seq data within this paper. For the P2 *Emx1-Eyfp* merge containing all cell types (Figure 1A), a resolution of 0.8 was used. For the P2 cell cycle regressed merge of *Emx1-Eyfp* and cortical datasets containing only V-SVZ lineages (Figure 1E), a resolution of 0.8 was used. For the E14, P2, P6/7 merge of *Emx1-Eyfp*-expressing RPs and NSCs, a resolution of 0.4 was used.

For the P2 *Emx1-Eyfp* (n2) dataset containing all cell types (Figure S1A), a resolution of 0.8 was used. For the previously published P2 *Emx1-Eyfp* (n1) dataset containing all cell types (Figure S1C), a resolution of 0.8 was used. For the P2 *Emx1-Eyfp* merge containing only cells that expressed *Emx1-Eyfp* > 1.5 (Figure S1D), a resolution of 0.8 was used. For the P2 *Emx1-Eyfp*/cortical merge containing V-SVZ lineages (Figure 1H), a resolution of 0.4 was used. For the E14 *Emx1-Eyfp* subset containing only RPs expressing *Emx1-Eyfp* > 1 (Figure S1E), a resolution of 0.8 was used. For the P6/7 *Emx1-Eyfp* merge containing all cells that expressed *Emx1-Eyfp* > 1.25 (Figures S1F-S1G), a resolution of 0.8 was used. For the previously published P2 cortical dataset containing all cell types (Figure S1H), a resolution of 0.8 was used.

For the E17/18 *Emx1-Eyfp*/cortex merged dataset containing only major V-SVZ cell lineages (Figure 2A), a resolution of 1.6 was used. For the E14, E17/18, P2, P6/7 merge of all cortically derived RPs and NSCs (Figure 2D), a resolution of 0.4 was used.

For the E18 cortical dataset containing all cell types (Figure S2A), a resolution of 2.4 was used. For the E17 *Emx1-Eyfp* dataset containing all cell types (Figure S2B), a resolution of 0.8 was used. For the P6/7 *Emx1-Eyfp*/cortical merge containing all cell types (Figure S2E), a resolution of 0.8 was used.

### Trajectory inference using transcriptomic data

Trajectory inference and pseudotime ordering of scRNA-seq data was performed using the Monocle 2 package (v2.20.0)^66^ in R as previously described.^4,27,64,67^ In short, the raw expression data for the cells of interest was extracted and merged. Following this, the data was normalized using Monocle’s size factor normalization method via the estimateSizeFactors function. PCA was then performed using the same 2,000 highly variable genes obtained from our computational pipeline, as described above, with slight modification if cell cycle regression was to be performed (see *Cell cycle regression*). Afterwards, the cells were projected into two-dimensional space and clusters were assigned using Monocle’s density peak clustering algorithm via the reduceDimension (max_components = 2, reduction_method = “tSNE”) and clusterCells functions in Monocle. Genes to be used for pseudotime ordering were obtained by identifying the top 1,000 differentially expressed genes between either clusters at the relevant resolution (Figure 1J) or timepoints (Figure 2H) using the differentialGeneTest function. Expression profiles were then reduced to two dimensions via the DDRTree algorithm using the reduceDimension function in Monocle (max_components = 2, method = “DDRTree”), and cells were then ordered using these genes to generate trajectories via the orderCells function. The function plot_cell_trajectory was subsequently used for trajectory visualization.

### Cell cycle regression

Cell cycle regression, in the context of clustering and visualizing scRNA-seq data (see Figure 1H), was performed by assigning cell cycle scores to the cells within the dataset based on their expression of cell cycle phase markers. Cell cycle scoring was performed using the CellCycleScoring function in the Seurat package. The ScaleData function was then used to generate a scaled expression matrix in which cell cycle-related heterogeneity was removed from the data by regressing out S and G2/M scores. All upstream and downstream steps were performed as previously discussed.

In order to regress cell cycle related genes from trajectories generated using Monocle, cell cycle associated genes used by Cyclone to assign cell cycle phase,^32^ along with S phase- and G2/M phase-related genes identified by Kowalczyk et al.^68^ and Tirosh et al.,^69^ were subtracted from the set of 1,000 ordering genes used to order trajectories in pseudotime, as described by Borrett and colleagues.^4^ Upstream and downstream steps were performed as described in the relevant section.

### Multiomic data analysis

snRNA-seq data was processed through the Seurat-based scRNA-seq pipeline described earlier. snATAC-seq data was analyzed using the ArchR R package (v1.0.3).^33^ In short, fragments output files generated by 10x Genomics Cell Ranger were read and processed using ArchR, excluding low-quality cells with <3000 fragments and a TSS enrichment score < 4. Cells were also removed based on high nucleosome ratio (>4) and blacklist ratio (>0.04) values. Genes were defined as a region containing the gene body as well as 5 kb upstream of the TSS and gene accessibility scores, which we used as a measure of the overall accessibility of a particular gene, were calculated using ArchR via the addGeneScoreMatrix function using default parameters. In short, fragments that mapped to a region 100 kb upstream and downstream of a given gene (unless this window overlapped with an adjacent gene, in which case it would be truncated accordingly) were used to calculate a gene score in a distance-dependent fashion, with distal fragments contributing less to accessibility calculations. Calculations were also normalized based on gene body size.

Dimensionality reduction was performed using an iterative LSI approach via the ArchR function addIterativeLSI (iterations = 2, clusterParams = list(resolution = c(0.2), sampleCells = 10000, n.start = 10), varFeatures = 25000, dimsToUse = 1:30). Clustering of the data, as implemented by the Seurat package ^30^, was performed across a range of resolutions (from 0.4 to 2.4) using the addClusters function (reducedDims = “IterativeLSI”, method = “Seurat”). Each dataset was analyzed by choosing the most conservative resolution in which cell types of known identity separated into distinct clusters as expected; resolution was only increased to interrogate cell heterogeneity or if two cell populations of known identity, based on the expression of canonical markers, were not separating into distinct clusters as expected. The data was projected into two-dimensional space as UMAP embeddings using the addUMAP function. Imputation, using the addImputeWeights function in ArchR with default parameters, was performed using Markov affinity-based graph imputation of cells (MAGIC)^70^ for clearer visualization of accessibility data. UMAPs and UMAP overlays, generated using the plotEmbedding function, were used to investigate gene expression and interpret cell clustering patterns.

Some clusters with unusually high fragment counts were considered to be low-quality cells and were removed. To identify cell types, we used the following marker genes: oligodendrocyte lineage cells expressed *Olig2* and *Sox10*; OPCs expressed *Pdgfra* and *Cspg4*; mature oligodendrocytes expressed *Mbp*, *Mog*, *Mag*, *Trf*, and *Opalin*; immature oligodendrocytes expressed *Fyn, Enpp6*, and *Gpr17*; both NSCs and astrocytes expressed *Slc1a3*, *Gfap*, and *Aldoc*; NSCs expressed *Veph1*, *Shroom3*, *Tspan18*, *Tfap2c*, and *Vnn1*; astrocytes expressed *Htra1*, *Aqp4*, and *Agt*; TAPs expressed *Ascl1*, *Egfr*, and *Gsx2*; olfactory bulb neuroblasts expressed *Dlx1*, *Dlx2*, *Sp8*, *Sp9*, *Gad1*, and *Gad2*; excitatory neurons expressed *Kitl*, *Ldb2*, *Neurod1*, *Satb2*, *Tbr1*, and *Unc5d*; microglia expressed *Cd68*, *Cx3cr1*, and *Aif1*; striatal interneurons expressed *Gad1*, *Ppp1r1b*, *Bcl11b*, and *Calb1*; cortical interneurons expressed *Sst*, *Pvalb*, and/or *Vip* in addition to *Gad1*; Cajal-Retzius cells expressed *Cacna2d2*, *Ebf3*, *Tmem163*, and *Trp73*; pericytes expressed *Cspg4*, *Pdgfrb*, *Des*, and *Rgs5*; endothelial cells expressed *Pecam1*, *Esam*, and *Plvap*; proliferating cells expressed *Ki67* and *Top2a*.

Merging and subsetting of the snRNA-seq data was performed as described above for scRNA-seq analyses. To subset snATAC-seq data, barcodes corresponding to the cells or cell types of interest were first obtained and the subsetArchRProject function was then used to remove all other cells; the data was then run through the pipeline as described above. To merge snATAC-seq data, an ArchRProject was first generated using all datasets to be merged. Afterwards, if relevant, barcodes corresponding to the cells or cell types of interest were subsetted out using the subsetArchRProject function and the data was run through the pipeline as described above.

For some downstream analyses, gene accessibility scores, motif accessibility z-scores, embeddings, and clustering, which were calculated using ArchR, were imported into Seurat/Signac.^71^ To do so, some code from the ArchRtoSignac R package^72^ was consulted and adapted.

### Batch correction multiomic data

Batch correction of multiomic data was performed using the Harmony package in R (v1.2.0).^31^ All batch correction of multiomic data was performed at the transcriptomic level. In brief, the RunHarmony function was used to reduce batch effects that manifested between cells originating from different datasets within a Seurat object. All multiomic datasets were assigned to the following batches, which largely corresponded to when each dataset was generated, and run through Harmony accordingly. Batch 1 consisted of the P7 V-SVZ dataset. Batch 2 consisted of the P6 V-SVZ dataset. Batch 3 consisted of the P2 V-SVZ (n1) dataset. Batch 4 consisted of the E14 cortex dataset. Batch 5 consisted of the E17 cortex and P2 V-SVZ (n2) datasets. The minimum number of Harmony iterations (either 1 or 2) required for the datasets to intermingle as expected were performed; the number of Harmony iterations used per dataset is described where relevant in the results section of this paper. After using RunHarmony, new UMAP embeddings were generated using the RunUMAP function, and the data was reclustered using the FindNeighbors and FindClusters functions at a range of resolutions.

### Resolutions used for multiomic data annotation and visualization

The following resolutions were used to visualize multiomic data within this paper. For the E14 cortical dataset containing V-SVZ lineages, clustered based on transcriptomic data (Figure 3A), a resolution of 0.8 was used. For the E14 cortical dataset containing V-SVZ lineages, clustered based on epigenetic data (Figure 3B), a resolution of 0.8 was used. For the E17 cortical dataset containing V-SVZ lineages, clustered based on transcriptomic data (Figure 3D), a resolution of 0.8 was used. For the E17 cortical dataset containing V-SVZ lineages, clustered based on epigenetic data (Figure 3F), a resolution of 0.4 was used. For the P2 V-SVZ merge containing V-SVZ lineage, clustered based on transcriptomic data (Figure 3G), a resolution of 0.8 was used. For the P2 V-SVZ merge containing V-SVZ lineage, clustered based on epigenetic data (Figure 3H), a resolution of 0.8 was used. For the P6/7 V-SVZ merge containing V-SVZ lineages, clustered based on transcriptomic data (Figure 3I), a resolution of 1.2 was used. For the P6/7 V-SVZ merge containing V-SVZ lineages, clustered based on epigenetic data (Figure 3J), a resolution of 0.4 was used.

For the E14 cortical dataset containing all cell types, clustered based on transcriptomic data (Figure S3A), a resolution of 1.2 was used. For the E17 cortical dataset containing all cell types, clustered based on transcriptomic data (Figure S3C), a resolution of 0.8 was used. For the P2 SVZ (n1) dataset containing all cell types, clustered based on transcriptomic data (Figure S3E), a resolution of 1.2 was used. For the P2 SVZ (n2) dataset containing all cell types, clustered based on transcriptomic data (Figure S3F), a resolution of 2.0 was used. For the P7 V-SVZ dataset containing all cell types (Figure S3G), clustered based on transcriptomic data, a resolution of 0.8 was used. For the previously published P6 V-SVZ dataset containing all cell types (Figure S3H), a resolution of 0.4 was used.

For the E14, E17, P2, P6/7 merge containing all cortical RPs and NSCs, clustered based on transcriptomic data (Figure S4A), a resolution of 0.4 was used. For the E14, E17, P2, P6/7 merge containing all cortical RPs and NSCs, clustered based on epigenetic data (Figure S4A), a resolution of 0.4 was used.

### Peak calling and motif analysis

Cells were grouped into pseudo-bulk replicates via the addGroupCoverages function in ArchR. Pseudo-bulk replicates were generated within either clusters at a resolution sufficient to distinguish between the cell types of interest or the annotated cell types themselves, depending on what was more appropriate. The default settings (minCells = 40, maxCells = 500, minReplicates = 2, maxReplicates = 5) were used. Peak calling was subsequently performed with MACS2 (v2.2.7.1)^73^ using the addReproduciblePeakSet function in ArchR.

Transcription factor binding motif inference was performed using the implementation of chromVAR (v1.24.0)^34^ in the ArchR package. A set of murine motifs were retrieved from the CIS-BP database^74^ using the addMotifAnnotations function. Z-scores of the bias-corrected deviations of binding motif accessibility were used as a measure of the enrichment or depletion of motifs within individual cells. Z-scores were used for subsequent pairwise testing of differential motif accessibility between cell populations (see *Differential accessibility statistical analysis* below). For binding motif visualization, position weight matrices were extracted from the dataset’s motif matrix, converted into position probability matrices, and plotted using the ggseqlogo (v0.1) package.^75^ Binding motifs included a pseudocount of 0.008.^34^

To identify accessible peaks containing specific transcription factor binding motifs (see Figure 4J), the ArchR getMatches function was used; the output (most pertinently the matches assay) was then examined to identify peaks containing the motifs of interest. Chromatin and peak accessibility were then visualized using the ArchR plotBrowserTrack function.

### Xenium panel design

Single-cell spatial transcriptomics was performed using a panel targeting 347 genes. 247 of these probes were from the mouse brain panel designed by 10x Genomics. A 100-probe custom add-on panel was generated to target the following genes: *Dll3*, *Nfix*, *Slc38a1*, *Tek*, *Bmx*, *Agt*, *Hbegf*, *Eps8*, *Gjb6*, *Lcat*, *Icam2*, *Vcam1*, *Cxcl12*, *Cxcr4*, *Jag1*, *Notch1*, *Clic6*, *Kcnj13*, *Aldoc*, *Hes5*, *Mt3*, *Slc1a3*, *Rgcc*, *Ttyh1*, *Gsta4*, *Grasp*, *Cdh5*, *Dll4*, *Tie1*, *Foxj1*, *Crybb1*, *Ecscr*, *Epas1*, *Npas1*, *Nrarp*, *Pcp4*, *Ptprn*, *Hes1*, *Clu*, *Il1r1*, *Enpp6*, *Fyn*, *Angpt2*, *Vegfa*, *Il6*, *Il1b*, *Osm*, *Il6st*, *Lifr*, *Eomes*, *Neurod1*, *AB124611*, *Cybb*, *Lgals3*, *Wfdc17*, *Cd14*, *Ly96*, *Myd88*, *Nfkb1*, *Tlr4*, *Cebpd*, *Cfn*, *Icam1*, *Lcn2*, *Bmp7*, *Clec11a*, *Lama1*, *Slc22a6*, *Aif1*, *Tmem119*, *Dlx1*, *Dlx2*, *Dlx5*, *Sp8*, *Sp9*, *P2ry12*, *Mdfi*, *Nes*, *Shroom3*, *Tfap2c*, *Thbs4*, *Tspan18*, *Veph1*, *Vnn1*, *Fam107a*, *Prex2*, *Mag*, *Mbp*, *Olig1*, *Olig2*, *Gpnmb*, *Igf1r*, *Ascl1*, *Egfr*, *Gsx2*, *Ki67*, *Flt1*, *Nr4a1*, *Emx1*, and *Nkx2-1*. All probes, from both predesigned and custom panels, are listed in Table S4. When designing the custom add-on panel, relevant scRNA-seq datasets were used to validate the target gene expression and to minimize the risk of optical crowding on a per-gene basis.

### Xenium tissue preparation & data processing

Brain tissue was collected from mice at E15, E18, and P2. Samples were fixed in 4% paraformaldehyde (Electron Microscopy Sciences) for 24 hours at 4°C and subsequently cryopreserved in 30% sucrose for 48 hours. Samples were embedded in O.C.T. (Tissue-Tek) and cryosectioned coronally onto Xenium slides (chemistry v1) at a thickness of 10 μm. Slides were then stored at −70°C until downstream steps.

All samples were processed as per the Xenium workflow for fixed frozen tissue as previously described.^28^ In brief, slides were incubated at 37°C for 1 minute and subsequently underwent a series of rehydrating washes in RNase-free PBS (Thermo Fisher), RNase-free water (Wisent), 100% ethanol (Sigma), and 70% ethanol (Sigma). Samples were then incubated in a decrosslinking buffer for 30 minutes at 80°C, followed by 10 minutes at 22°C. Following three PBS-T washes, tissue was incubated in a probe solution for 16-24 hours at 50°C. For all datasets, the solution contained the same set of 347 gene targeting probes (see Table S4) in TE buffer (Fisher). Nonspecific probe binding was next washed off via PBS-T, a 30-minute incubation in post-hybridization buffer at 37°C, and additional PBS-T washes.

Ligation of the hybridized probes was performed by incubating the samples in ligation enzymes for 2 hours at 37°C. After a series of PBS-T washes, the ligated probes were amplified via rolling circle amplification; to achieve this, samples were incubated with amplification enzyme for 2 hours at 30°C. Sections were then washed twice in TE buffer.

Autofluorescence quenching and nuclei staining steps were carried out next. Sections were washed in PBS, incubated for 10 minutes in a reducing agent, washed in 70% and 100% ethanol, and incubated in Xenium autofluorescence solution for 10 minutes. Following this, sections were washed three times in 100% ethanol, dried at 37°C for 5 minutes, and rehydrated via subsequent washes in PBS and PBS-T. Sections were then incubated in a nuclear staining buffer for 1 minute and washed in PBS-T.

Following this, the sample was loaded into the Xenium analyzer instrument (software v2.0-3.1, depending on the sample) and subjected to multiple cycles of reagent application, probe hybridization, imaging, and probe removal. Initial pre-processing of captured Z-stack images was performed using the Xenium onboard analysis pipeline (v2.0-3.1, depending on the sample), as outlined extensively by Janesick and colleagues.^52^ In brief, our custom Xenium codebook was used to decode puncta into transcripts, with each codeword assigned to genes in the gene panel. Quality scores (Q-Scores) were assigned to transcripts based on maximum likelihood codewords, and negative controls were used to ensure accurate calibration. On-instrument nuclei segmentation was performed based on DAPI morphology, with each segmented nuclei assigned a unique cell ID.

All data were subsequently run through 10x Genomics Xenium Ranger v3 software (xeniumranger resegment function) with settings as recommended by the manufacturer to assign transcripts to the closest nucleus within a maximum distance of 2 µm.

### Xenium data analysis

Xenium data were processed using a previously described computational pipeline.^25,26,28^ Xenium data files were first visualized in Xenium Explorer (v1.3-4). Regions of interest (ROIs) were defined for each section. Across all timepoints, these ROIs encompassed the dorsal V-SVZ, some more ventral regions of the V-SVZ, and cortical tissue dorsal to the lateral ventricles. The cell IDs corresponding to each ROI were then imported into R alongside transcript count, feature, and coordinate data via the Seurat package (v5.2.0) to generate Seurat objects containing expression data corresponding to the regions of interest per tissue section. Low quality cells were identified based on the mean number of detected genes and total transcript counts, and cells greater than +/- 2.5 standard deviations from the mean were filtered out.

Filtered data were transformed using SCTransform and dimensionality reduction was performed using PCA based on highly variable genes. This was subsequently used to construct a shared nearest neighbor (SNN) graph using the FindNeighbors function in Seurat. The data was then clustered using the FindClusters function across a range of resolutions (0.4 to 2.4) and visualized in two-dimensional space using UMAP embeddings. Spatial plots and UMAPs, generated using the ImageDimPlot, FeaturePlots, and ImageFeaturePlots functions, were used to investigate gene expression and interpret cell clustering patterns.

Each section was initially processed separately. Afterwards, data corresponding to different sections were merged together in one of two ways. For merged datasets of the same timepoint (e.g. all E15 sections), data was integrated using the Seurat SelectIntegrationFeatures, FindIntegrationAnchors, and IntegrateData functions. For the dataset containing data from all three Xenium timepoints, merging was performed using the Seurat merge function, so as to avoid the loss of biologically significant differences between the timepoints. Where necessary for improved resolution of certain cell types, the Seurat subset function was used on merged datasets to subset out clusters of interest. Normalization, transformation, PCA, and clustering was performed on each merged dataset and/or subset as previously described.

Cell types were identified based on expression of canonical markers and positional information. We used the following gene expression definitions: oligodendrocyte lineage cells expressed *Olig1*, *Olig2*, and *Sox10*; OPCs expressed *Pdgfra* and *Cspg4*; immature oligodendrocytes expressed *Enpp6*, and *Gpr17*; mature oligodendrocytes expressed *Mbp* and *Mag*; NSCs expressed *Tspan18*, *Veph1*, *Shroom3*, *Tfap2c*, *Vnn1*, and *Thbs4*; astrocytes expressed *Hbegf*, *Agt*, *Lcat*, and *Gjb6*; TAPs expressed *Ascl1*, *Egfr*, and *Gsx2*; olfactory bulb neuroblasts expressed *Dlx1*, *Dlx2*, *Dlx5*, *Sp8*, and *Sp9*; cortical excitatory neurons expressed *Neurod6* and *Slc17a7* as well as *Bcl11b*, *Cux2*, *Fezf2*, and/or *Satb2*; IPs expressed *Eomes* and only lowly expressed *Neurod1*; late IPs expressed *Eomes* and *Neurod1*; microglia expressed *Aif1*, *Trem2*, and *Cd68*; striatal interneurons expressed *Ppp1r1b*, *Calb1*, and *Bcl11b*; cortical interneurons expressed *Sst*, *Pvalb*, and/or *Vip* in addition to *Gad1*; septal interneurons expressed *Gad1*, *Sp8*, *Cacna2d2*, and *Calb2*; endothelial cells expressed *Pecam1*, *Acvrl1*, *Adgrl4*, and *Car4*; ependymal cells expressed *Foxj1*; choroid plexus expressed *Clic6*; proliferating cells expressed *Ki67*. Cells in the P2 dataset that expressed both NSC and TAP markers and localized relatively far from the ventricle were termed “corner precursors”.

Each dataset was analyzed by choosing the most conservative resolution; resolution was only increased to interrogate cell heterogeneity or if two cell populations of known identity, based on the expression of canonical markers, were not separating into distinct clusters as expected.

### Resolutions used for Xenium data annotation and visualization

The following resolutions were used to annotate and visualize Xenium data within this paper. For the E18 dataset containing all cell types (Figure 5B), cell type annotations were based on clustering at a resolution of 1.2. For the E18 subset containing major precursor populations (Figure 5I), cell type annotations were based on clustering at a resolution of 2.0.

For the E15 dataset containing all cell types (Figure 6F), cell type annotations were based on clustering at a resolution of 0.4.

For the P2 dataset containing all cell types (Figure 7A), cell type annotations were based on clustering at a resolution of 2.0. For the E15, E18, P2 merged dataset containing major precursor populations (Figures 7I and S7J), a resolution of 1.2 was used.

### Xenium Explorer visualizations

Xenium Explorer (v4) was used to visualize expression of individual mRNAs on tissue sections. To do so, cells, nuclear boundaries, and detected transcripts were overlaid on top of DAPI staining. For cell type visualizations, Seurat-based cluster assignments at the relevant resolution were imported into Xenium Explorer and clusters were colored based on their cell type annotations.

## Quantification and Statistical Analysis

### Differential gene expression analysis

Differential gene expression analysis of scRNA-seq data were performed using the FindMarkers function in Seurat. Expression levels were tested using a Wilcoxon Rank Sum Test. A Bonferroni-adjusted p-value smaller than 0.05 (p_adj_ < 0.05) was considered statistically significant, and data was further filtered to only include genes with an average fold-change of greater than 1.3 (FC > 1.3), expression in at least 10% of cells within a given cluster, and a difference in the percent of cells expressing the gene between the groups being compared of at least 10%.

Differential gene expression analysis of Xenium data was performed using SCT-normalized expression data. This was performed using the PrepSCTFindMarkers function followed by the FindMarkers function in Seurat. Expression levels were tested using a Wilcoxon Rank Sum Test. A Bonferroni-adjusted p-value smaller than 0.05 (p_adj_ < 0.05) was considered statistically significant.

### Differential accessibility analysis

Bias-matched differential accessibility testing (of genes and transcription factor binding motifs) of multiomic data was performed using the ArchR getMarkerFeatures function. Cells assigned to the first group were compared against a second group, which was selected based on TSS enrichment values, log-normalized fragment counts, and log-normalized transcript counts in order to reduce potential bias. Expression levels were tested using a Wilcoxon Rank Sum Test. A false detection rate smaller than 0.05 (FDR < 0.05) combined with a log fold-change greater than 0.5 (log2FC > 0.5) was considered statistically significant.

### Gene signature analysis

Scaled expression or accessibility of multiple genes was calculated using the AddModuleScore function in R. Module scores were calculated using the log-normalized gene expression or gene accessibility scores. To calculate the overall expression/accessibility of the postnatal NSC signature, the following modified signature from Borrett et al.^4^ was used: *Sfrp1*, *Veph1*, *Tnc*, *Thbs4*, *Vnn1*, *Cpe*, *Igfbp5*, *Sparc*, *Mdfi*, and *Ccdc80.* Relative to the original signature, genes were removed if they were (i) non-specific to NSCs, (ii) sparsely expressed, or (iii) broadly expressed in cortical RP/NSCs.

The followed embryonic RP signature was defined based on differential expression analyses comparing E14 RPs and P6/7 NSCs (Table S2): *Col2a1*, *Neurog2*, *Car14*, *Sema5a*, *Bcl11b*, *Basp1*, *Neurod2*, *Eomes*, *Neurod6*, *Sdc1*, *Trp53i11*, *Jpt2*, *Nme4*, *Dmrta1*, and *Myo1b*.

### Pearson correlation analysis

Two types of correlation analyses were performed within this study. In the first, Pearson correlation analysis was performed by averaging the expression of a given transcript within two clusters/cell types of interest to obtain an averaged transcriptome for each group. Cell cycle associated genes used by Cyclone to assign cell cycle phase,^32^ along with S phase- and G2/M phase-related genes identified by Kowalczyk et al.^68^ and Tirosh et al.,^69^ were excluded from this analysis. Normalized expression values were then correlated using the cor.test function in R. For each comparison, the Pearson correlation coefficient (r) value is provided in the figure, figure legend, and results text.

In the second type of correlation analysis, single-cell correlation analysis, transcriptomes or epigenomes of individual cells were compared against averaged transcriptomes/epigenomes, once more using the cor.test function in R. For transcriptomic data, correlations were performed using log-normalized gene expression data; for epigenetic data, correlations were performed using log-normalized gene accessibility scores as calculated using the ArchR package. Individual cells were then plotted as a scatter plot, where placement along each axes represents the difference between the cell’s correlation to one averaged transcriptome/epigenome versus another.

### Xenium-based quantification & statistical analysis

All Xenium quantifications were performed using the E18 dataset, which contained 1 section each from 4 CD1 E18 brains collected across two litters. Roughly equivalent ROIs were used for each section; these ROIs all contained the entirety of the dorsal wall of the V-SVZ, the dorsal regions of the lateral and septal walls, the intermediate zone, and part of the cortical plate.

To quantify highly *Ki67*-expressing cells within dorsolateral and dorsomedial RP/NSCs in the E18 cortex (Figure 5N), we extracted the barcodes of cells that detectibly expressed 4 or more *Ki67* mRNAs. These *Ki67*-expressing cells were then cross-referenced with dorsolateral and dorsomedial RP/NSCs, as defined by the E18 subset (see Figure 5I), and the data was expressed as the proportions of cells within each RP/NSC population that expressed 4+ *Ki67* mRNAs. Statistical significance was determined using a paired parametric t-test, wherein values corresponding to the same section were paired, with p < 0.05 considered to be statistically significant.

To quantify the distribution of cortically derived TAPs within the E18 cortex (see Figures 6A and 6B), we used TAPs as identified by the E18 subset (Figure 5I). We defined TAPs of cortical origin as those expressing any amount of either *Emx1*, *Fezf2*, or *Neurod6*, or at least two *Tfap2c* mRNAs. The dorsal V-SVZ was then divided into dorsolateral and dorsomedial regions of equal length, and the number of cortically derived cells dorsal to these regions were quantified. TAPs were excluded if they were medial or lateral to the dorsal V-SVZ. The data was expressed as the proportion of cortical TAPs across both regions that were adjacent to either the dorsolateral or dorsomedial V-SVZ. Statistical significance was determined using a paired parametric t-test, wherein values corresponding to the same section were paired, with p < 0.05 considered to be statistically significant.

To quantify the identities of RP/NSCs adjacent to TAPs at E18 (see Figures 6C and 6D), we used TAPs, dorsolateral RP/NSCs, and dorsomedial RP/NSCs as defined by the E18 subset (Figure 5I). Cell type annotations were imported into and visualized in Xenium Explorer (v4) and manually quantified using Photoshop (Adobe). A TAP was considered adjacent to an RP/NSC if it was either (i) directly touching an RP/NSC, (ii) in close proximity to an RP/NSC without any cells directly between them, or (iii) directly touching another TAP considered to be adjacent to an RP/NSC. TAPs were excluded if they were medial or lateral to the dorsal V-SVZ. The data was expressed as the proportion of total RP/NSC-adjacent TAPs that were adjacent to either dorsolateral RP/NSCs, dorsomedial RP/NSCs, or at least one of each. Statistical significance was determined using a paired parametric t-test, and values corresponding to the same section were paired, with p < 0.05 considered to be statistically significant.

To quantify the distribution of RPs, transition precursors, and NSCs (see Figure 7L), we defined cells as per the E15/E18/P2 precursor merge (Figure 7I) and visualized E18 cells that clustered with cortical RPs (clusters 0 and 11), cortical transition precursors (cluster 7), and cortical NSCs (cluster 15) within the E18 cortex. As per the TAP localization analysis, the dorsal V-SVZ was then divided into dorsolateral and dorsomedial regions of equal length, and the number of cortically derived cells dorsal to these regions were quantified. Precursors were excluded if they were medial or lateral to the dorsal V-SVZ. The precursors present within the dorsolateral and dorsomedial V-SVZ were quantified using Illustrator (Adobe) and used to calculate the proportion of precursors per group that were localized to each region. To account for disparity in precursor abundance between the dorsomedial and dorsolateral V-SVZ, dorsolateral and dorsomedial proportions were divided by the overall distribution of precursors (clusters 0, 7, 11, 15) per section. In the resulting normalized data, a value of 1 represents a cluster distribution that closely mirrors the overall distribution of precursors; non-zero values represent either an over- or under-representation of precursors. Statistical significance between differentially distributed precursors within each cluster was determined using a two-way ANOVA with Bonferroni post-hoc comparisons, with p < 0.05 considered to be statistically significant.

For all plots, error bars indicate standard deviation (SD). Statistical analyses were performed using Prism 10 (GraphPad). Statistics used for computational analyses are described in the relevant sections.

**Table S1. *Genes differentially accessible between E14 cortical RPs and cortical precursors at E17, P2, and P6/7.* (Related to Figures 4 and S4.)** Shown are genes significantly differentially accessible between E14 cortical RPs and E17 RP/NSCs, P2 NSCs, and P6/7 NSCs. The comparison was performed between timepoints in the E14, E17, P2, and P6/7 RP/NSC merge shown in Figures 4A-4B. Genes were considered to be significantly enriched if their log fold-change was greater than 0.5 (log2FC > 0.5) and the gene had a false discovery rate of less than 0.05 (FDR < 0.05). Genes mentioned in the results are highlighted.

**Table S2. *Genes differentially expressed between E14 cortical RPs and P6/7 cortical NSCs.* (Related to Figures 4 and S4.)** Shown are genes significantly differentially expressed between E14 cortical RPs and P6/7 cortical NSCs. The comparison was performed between the E14 and P6/7 timepoints in Figures 2D-2E. Genes were considered to be significantly enriched if their average expression was at least 1.3-fold upregulated in one group versus the other (avg. fold-change > 1.3), the gene had a Bonferroni-adjusted p-value of less than 0.05 (p adj < 0.05), the gene was expressed in at least 10% of precursors in either group (% E14 RPs > 10% or % P6/7 NSCs > 10%), and there was a 10% difference in the proportion of precursors expressing the gene between groups (% diff > 10%). Highlighting indicates RP signature genes.

**Table S3. *Transcription factor binding motifs differentially accessible between E14 cortical radial precursors and P6/7 cortical neural stem cells*. (Related to Figures 4 and S4.)** Shown are genes significantly differentially accessible between E14 cortical RPs and P6/7 cortical NSCs. The comparison was performed between the E14 and P6/7 timepoints in Figures 4A-4B. Genes were considered to be significantly enriched if the gene had a false discovery rate of less than 0.05 (FDR < 0.05). Included in the table is an “Index” column, in which a unique identifier has been appended to the name of the transcription factor; this is to distinguish between motifs in cases where a transcription factor has multiple. Genes mentioned in the results are highlighted.

**Table S4. *Probes used for Xenium single-cell spatial transcriptomics.* (Related to Figures 5-7 and S5-S7.)** Shown are the genes included in the 100-gene Xenium custom add-on probeset, as well as the 247-gene Xenium pre-designed mouse brain panel, which were used for all Xenium experiments. Included are Ensembl IDs for all genes used, as well as the number of probes used for each gene in the custom add-on; the probe number was tuned to allow for optimal detection while avoiding optical crowding.

**Table S5. *Genes differentially detected between dorsolateral and dorsomedial RP/NSCs at E18.* (Related to Figures 5 and S5.)** Shown are Xenium probes differentially detected in a comparison between dorsolateral and dorsomedial RP/NSCs, as defined in the E18 subset shown in Figure 5I. Probes were considered to be significantly enriched if the gene had a Bonferroni-adjusted p-value of less than 0.05 (p adj < 0.05) and the gene was expressed in at least 5% of precursors in either group (% dorsolateral > 5% or % dorsomedial > 5%). Also shown are the average fold-change (avg. fold-change) and the log-normalized fold-change (avg. log2FC). Genes mentioned in the results are highlighted.

