## Supplemental Figures and Legends for "Single-cell multiomic approaches define a gradual, spatially-regulated epigenetic and transcriptional transition from embryonic to adult neural stem cells"

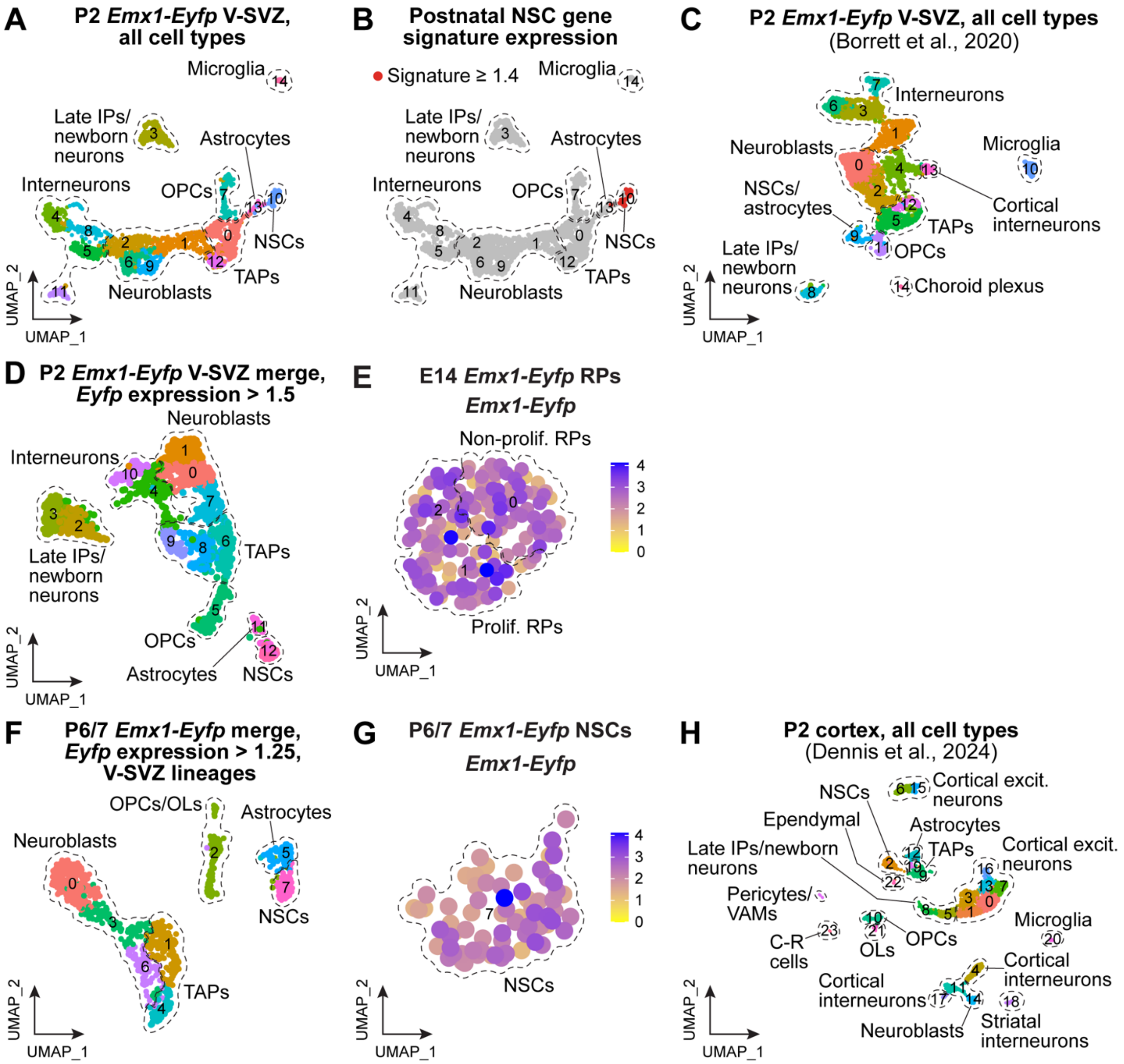

**Figure S1.** ***scRNA-seq datasets used to transcriptionally-define P2 cortical V-SVZ cell lineage cells.*** **(Related to Figure 1). (A, B)** Uniform manifold approximations and projections (UMAPs) of transcriptomes from a P2 *Emx1-Eyfp* ventricular-subventricular zone (V-SVZ) scRNA-seq dataset. (A) is annotated for cell types (indicated by hatched lines), with different colors denoting transcriptionally distinct clusters. (B) shows cells expressing a postnatal NSC gene signature described in Borrett et al.,[1] with modifications. Red denotes cells that express the gene signature at a level of 1.4 or above (see Experimental Methods for details). **(C)** UMAP of transcriptomes from a previously published scRNA-seq dataset (Borrett et al.,[1] GEO: GSE152281) collected from the *Emx1-Eyfp* V-SVZ at P2 and reprocessed through our pipeline. Transcriptionally distinct clusters are color-coded and cell types are annotated (indicated by hatched lines). **(D)** The merged P2 *Emx1-Eyfp* V-SVZ dataset in (Figure 1A) was subsetted to include only cells with normalized *Emx1-Eyfp* expression levels above 1.5 (as in Figure 1D), reanalyzed via our pipeline, and batch-corrected via two iterations of Harmony. The UMAP shows the resultant cluster diagram with cell types annotated (indicated by hatched lines) and transcriptionally distinct clusters color-coded. **(E)** UMAP of *Emx1-Eyfp* cortical RP transcriptomes from previously-published scRNA-seq datasets (Borrett et al.,[1] GEO: GSE152281) reprocessed through the pipeline to only include cells with normalized *Emx1-Eyfp* mRNA expression above 1. The UMAP shows *Emx1-Eyfp* expression levels, color-coded as per the adjacent key. The hatched lines and numbers denote transcriptionally-distinct clusters. **(F)** V-SVZ neural cell lineage transcriptomes from previously published P6/7 *Emx1-Eyfp* cortex scRNA-seq datasets (Borrett et al.,[1] GEO: GSE152281) reprocessed through the pipeline to only include cells with normalized *Emx1-Eyfp* mRNA expression above 1.25. Transcriptionally distinct clusters are color-coded and cell types are annotated (indicated by hatched lines). **(G)** NSC transcriptomes from (F; cluster 7), overlaid for *Emx1-Eyfp* mRNA expression, color-coded as per the adjacent key. **(H)** UMAP of transcriptomes from a previously published P2 cortex scRNA-seq dataset (Dennis et al.,[2] GEO: GSE255405), reanalyzed via the pipeline showing color-coded transcriptionally distinct clusters and cell type annotations (indicated by hatched lines). **Abbreviations:** C-R, Cajal-Retzius. Excit., excitatory. IPs, intermediate progenitors. NSCs, neural stem cells. OLs, oligodendrocytes. OPCs, oligodendrocyte precursor cells. TAPs, transit-amplifying precursors. VAMs, vascular associated mesenchymal cells. V-SVZ, ventricular-subventricular zone.

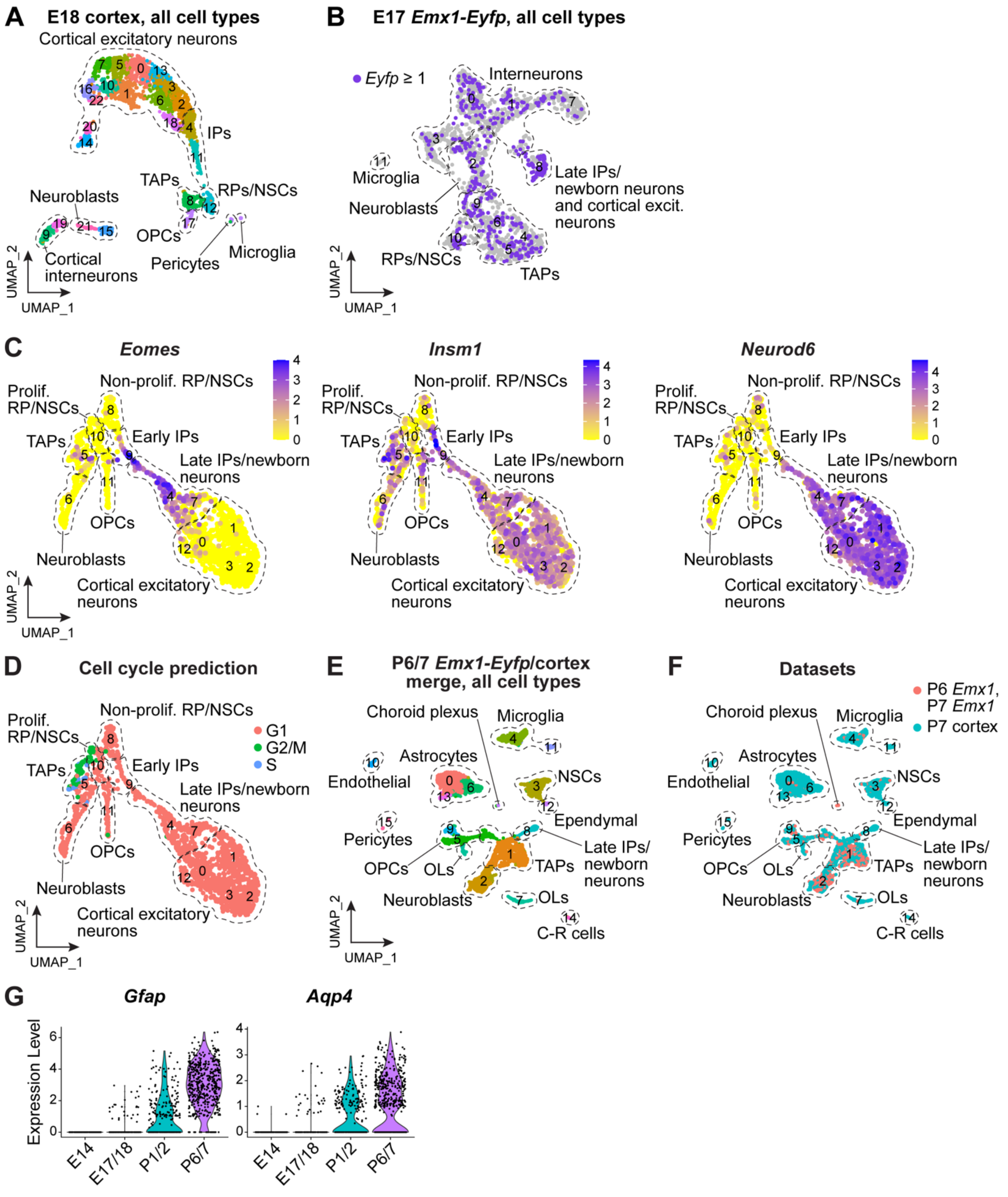


**Figure S2.** ***scRNA-seq datasets used to transcriptionally-define E17/18 cortical V-SVZ cell lineage cells.*** **(Related to Figure 2). (A)** UMAP of transcriptomes from an E18 cortex scRNA-seq dataset showing cell type annotations (indicated by hatched lines) and transcriptionally distinct cell clusters, color-coded. **(B)** UMAP of transcriptomes from a previously published

*Emx1-Eyfp* E17 V-SVZ scRNA-seq dataset (Borrett et al.,[1] GEO: GSE152281) reprocessed through the pipeline. Cell types are annotated (indicated by hatched lines), and cells expressing *Emx1-Eyfp* mRNA at normalized levels greater than 1 are color-coded purple. Cells that do not meet this threshold are gray. **(C)** Gene expression overlays on UMAPs of the merged E17/18 V-SVZ dataset in Figure 2A showing normalized expression of the IP marker mRNAs *Eomes* and *Insm1*, as well as *Neurod6*, which is highly enriched in late IPs/newborn neurons, color-coded as per the adjacent keys. Cell types are also annotated (indicated by hatched lines). **(D)** UMAP of the merged E17/18 V-SVZ dataset in Figure 2A showing cell cycle status as predicted using Cyclone. Cell cycle stages are color-coded as per the adjacent key. **(E, F)** Transcriptomes from a previously-published P7 cortex scRNA-seq dataset (Dennis et al.,[2] GEO: GSE255405) were reprocessed through the pipeline, merged with *Emx1-Eyfp*-expressing transcriptomes from the P6/7 *Emx1-Eyfp* V-SVZ (Figure S1F), and batch-corrected using one iteration of Harmony. The UMAP in (E) shows color-coded, transcriptionally-distinct clusters, while (F) shows the datasets of origin, color-coded as per the adjacent key. In both panels, cell types are annotated (indicated with hatched lines). **(G)** Violin plots showing normalized expression of the NSC marker mRNAs *Gfap* and *Aqp4* within E14, E17/18, P2 and P6/7 cortical precursors, all from the dataset in Figure 2D, separated by timepoints. Each dot represents expression levels in an individual cell. **Abbreviations:** C-R, Cajal-Retzius. INs, interneurons. IPs, intermediate progenitors. NSCs, neural stem cells. OLs, oligodendrocytes. OPCs, oligodendrocyte precursor cells. Prolif., proliferating. RPs, radial precursors. TAPs, transit-amplifying precursors.

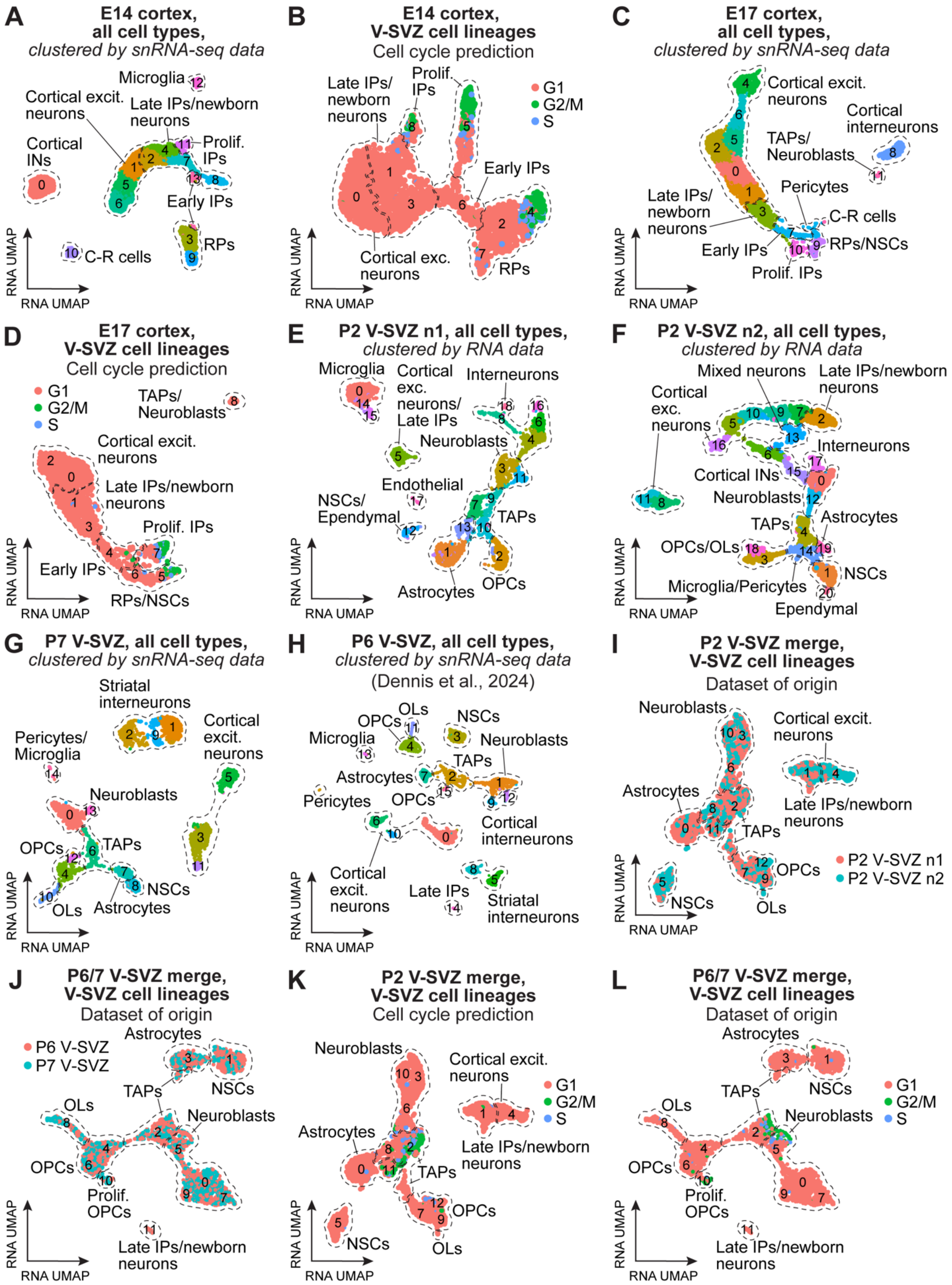
A
E14 cortex, all cell types, clustered by snRNA-seq data
B
E14 cortex, V-SVZ cell lineages
Cell cycle prediction
C
E17 cortex, all cell types, clustered by snRNA-seq data
D
E17 cortex, V-SVZ cell lineages
Cell cycle prediction
E
P2 V-SVZ n1, all cell types, clustered by RNA data
F
P2 V-SVZ n2, all cell types, clustered by RNA data
G
P7 V-SVZ, all cell types, clustered by snRNA-seq data
H
P6 V-SVZ, all cell types, clustered by snRNA-seq data (Dennis et al., 2024)
I
P2 V-SVZ merge, V-SVZ cell lineages
Dataset of origin
J
P6/7 V-SVZ merge, V-SVZ cell lineages
Dataset of origin
K
P2 V-SVZ merge, V-SVZ cell lineages
Cell cycle prediction
L
P6/7 V-SVZ merge, V-SVZ cell lineages
Dataset of origin
G1
G2/M
S
P2 V-SVZ n1
P2 V-SVZ n2
P6 V-SVZ
P7 V-SVZ
RNA UMAP

**Figure S3.** ***Paired single nucleus RNA-seq and ATAC-seq (multiomic) datasets used to define V-SVZ cell lineage cells from E14 to P6/7 transcriptomically and epigenetically.*** **(Related to Figure 3). (A)** UMAP showing E14 cortex single nucleus multiomic data clustered on the basis of the transcriptional data. Transcriptionally-distinct clusters are colored and numbered. Cell types are annotated based upon marker gene expression (indicated by hatched lines). **(B)** UMAP of the E14 cortex V-SVZ neural cell lineage, subsetted from the dataset shown in (A) and reclustered on the basis of the transcriptional data (also see Figure 3A). The UMAP is annotated for cell types (indicated by hatched lines) and shows cell cycle states as predicted by Cyclone and color-coded as per the adjacent key. **(C)** UMAP showing the E17 cortex single nucleus multiomic dataset clustered on the basis of transcriptional data. Transcriptionally-distinct clusters are colored and numbered. Cell types are annotated based upon marker gene expression (indicated by hatched lines). **(D)** UMAP of the E17 cortex V-SVZ neural cell lineage subsetted from the dataset in (C) and clustered on the basis of the transcriptional data (also see Figure 3D). The UMAP is annotated for cell types (indicated by hatched lines) and shows cell cycle states as predicted by Cyclone and color-coded as per the adjacent key. **(E, F)** UMAPs of two different P2 V-SVZ single nucleus multiomic datasets (n1 and n2), clustered on the basis of the transcriptional data and annotated for cell types identified by marker genes (indicated by hatched lines). Transcriptionally-distinct clusters are colored and numbered. **(G)** UMAP showing the P7 V-SVZ single nucleus multiomic run, clustered on the basis of the transcriptional data and annotated for cell types identified by marker genes (indicated by hatched lines). Transcriptionally-distinct clusters are colored and numbered. **(H)** UMAP showing P6 V-SVZ single nucleus transcriptomes from a previously published multiomic analysis (Dennis et al.,[2] GEO: GSE255405). The data were reprocessed through our pipeline, clustered on the basis of the transcriptional data, and annotated for cell types identified by marker genes (indicated by hatched lines). Transcriptionally-distinct clusters are colored and numbered. **(I)** UMAP showing a merger of the P2 V-SVZ neural cell lineage multiomic datasets in (E and F), clustered on the basis of the transcriptional data (also see Figure 3G). The UMAP shows datasets of origin, color-coded as per the adjacent key and cell type annotations as in Figure 3G (indicated by hatched lines). **(J)** UMAP showing a merger of the P6/7 V-SVZ neural cell lineage multiomic datasets in (G and H) after one round of batch correction, clustered on the basis of the transcriptional data (also see Figure 3I). The UMAP shows the datasets of origin, color-coded as per the adjacent key and cell type annotations as in Figure 3I. **(K)** UMAP of the merged P2 V-SVZ cell lineage multiomic dataset shown in (I) and Figure 3G, clustered on the basis of the transcriptional data and annotated for cell types (indicated by hatched lines). Also shown are cell cycle states as predicted by Cyclone and color-coded as per the adjacent key. **(L)** UMAP of the merged P6/7 V-SVZ cell lineage multiomic dataset shown in (J) and Figure 3I, clustered on the basis of the transcriptional data, and annotated for cell types (indicated by hatched lines). Also shown are cell cycle states as predicted by Cyclone and color-coded as per the adjacent key. **Abbreviations:** C-R, Cajal-Retzius. Excit., excitatory. IPs, intermediate progenitors. NSCs, neural stem cell. OLs, oligodendrocytes. OPCs, oligodendrocyte precursor cells. Prolif., proliferative. TAPs, transit-amplifying precursors. RPs, radial precursors.

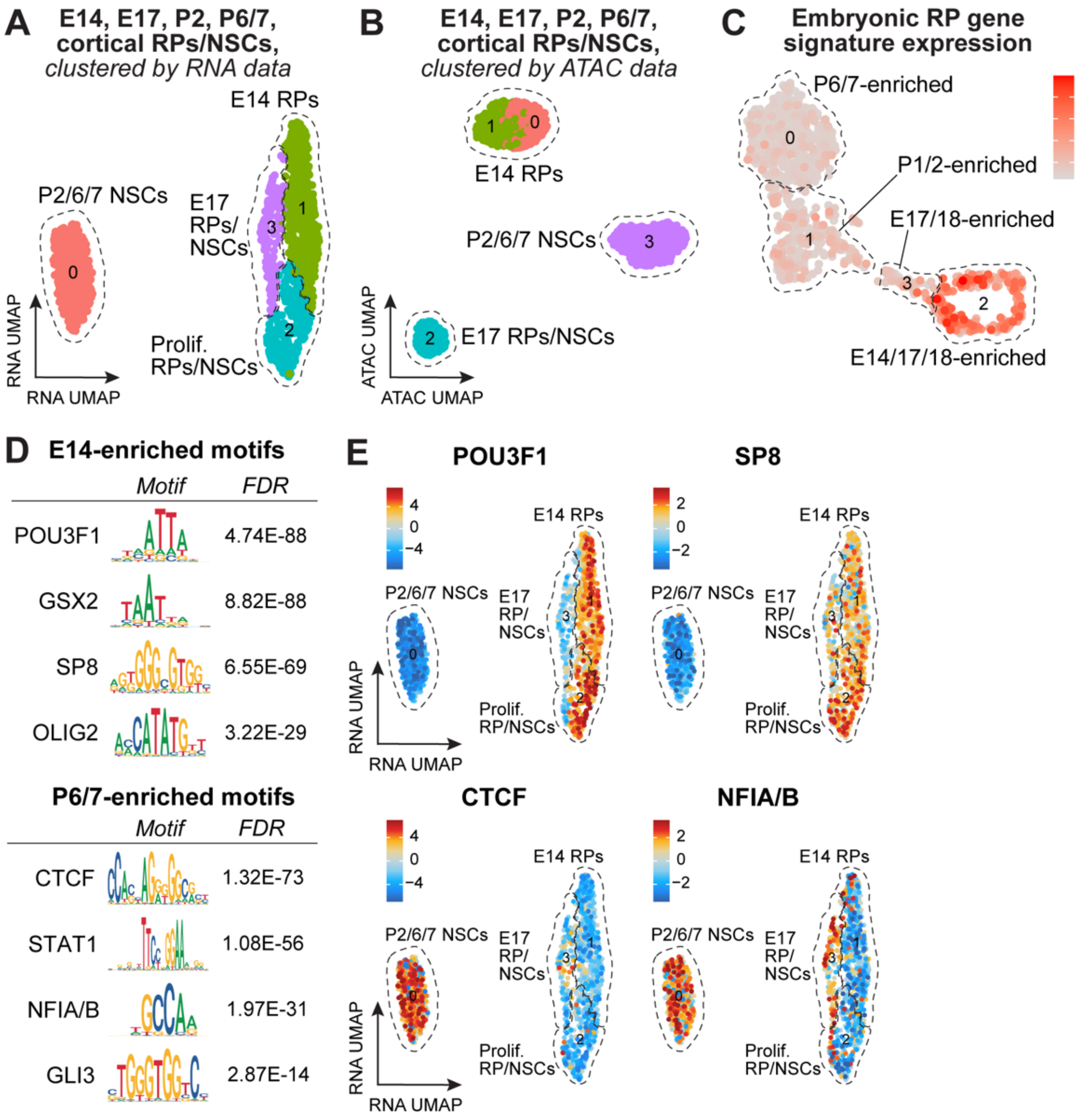


**Figure S4.** ***Comparison of cortical precursors from E14 to P6/7 using single nucleus multiomics.*** **(Related to Figure 4). (A)** UMAP of cortical precursor transcriptomes merged from the E14, E17, P2, and P6/7 multiomic datasets (shown in Figures 3A, 3D, 3G, and 3I) after one iteration of Harmony, clustered on the basis of the transcriptional data (also shown in Figure 4A). Transcriptionally-distinct clusters are color-coded and numbered, and cell types are annotated (indicated by hatched lines). **(B)** UMAP of the merged E14 to P6/7 cortical precursor multiomics dataset shown in (A), clustered on the basis of the paired snATAC-seq epigenomic data and annotated for cell types. Epigenetically-distinct clusters are color-coded and numbered. **(C)** UMAP overlay of a 15 gene embryonic RP signature defined by performing a differential gene expression comparison between E14 RPs and P6/7 NSCs (Table S2). The gene signature is visualized as a UMAP overlay on the merged E14 to P6/7 cortical precursor scRNA-seq dataset shown in Figure 2D. Expression levels are color-coded as per the adjacent key. **(D)** Logos and FDR values for select transcription factor binding motifs identified as being differentially accessible in a comparison of E14 versus P6/7 cortical precursor epigenomes (as shown in B) (Table S3). **(E)** UMAP of merged E14-P6/7 cortical precursors clustered on the transcriptional data as in (A) showing accessibility enrichment z-scores for selected transcription factor binding motifs identified as being differentially accessible in a comparison of E14 versus P6/7 cortical precursor epigenomes (those shown in B) (Table S3), color-coded as per the adjacent keys. The motifs used in this analysis are shown in (D). **Abbreviations:** NSCs, neural stem cells. RPs, radial precursors.

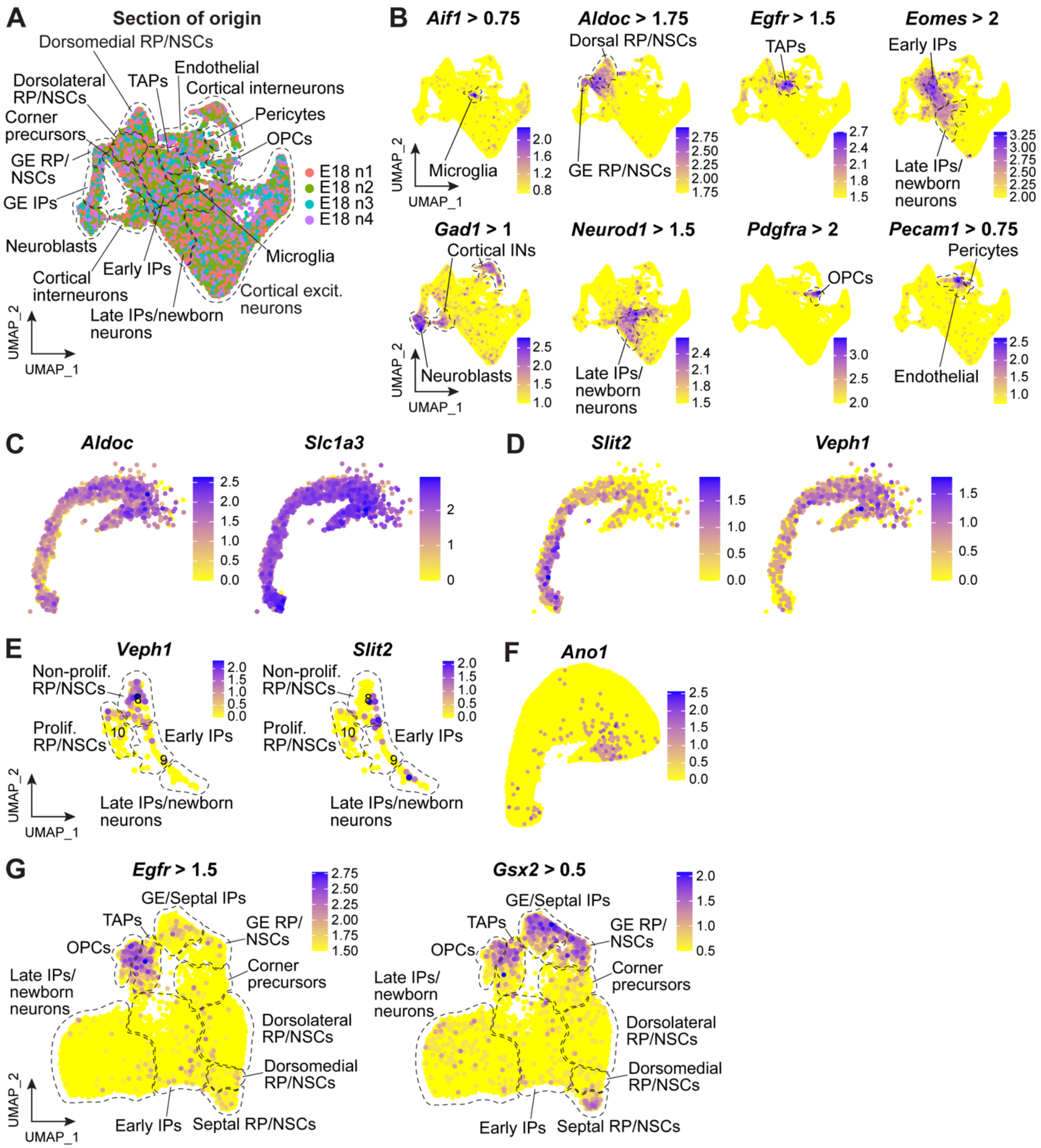


**Figure S5.** ***Characterization of E17/18 RP/NSCs and TAPs as detected using Xenium single-cell spatial transcriptomics*****. (Related to Figure 5). (A)** Xenium single-cell spatial transcriptomics was performed on coronal E18 cortical sections from 4 independent animals (1 each). Shown is a UMAP of the resultant merged transcriptomes from all 4 sections/animals, colored by dataset of origin, as per the adjacent key. Cell types are annotated (outlined by black lines), as in the UMAP shown in Figure 5B. Colors correspond to sections/animals. **(B)** UMAPs as in (A) showing thresholded expression of marker genes that distinguish microglia (*Aif1*), RP/NSCs (*Aldoc*), TAPs (*Egfr*), IPs (*Eomes*), inhibitory neuroblasts and neurons (*Gad1*), late IPs/newborn cortical neurons (*Neurod1*), OPCs (*Pdgfra*), and endothelial cells (*Pecam1*). Expression thresholds are indicated above each UMAP. **(C, D)** Spatial plots of the E18 RP/NSCs as defined in Figure 5J, overlaid for expression of mRNAs enriched (C) in all RP/NSCs (*Aldoc*, *Slc1a3*), or (D) in dorsolateral (*Veph1*) versus dorsomedial (*Slit2*) RP/NSCs. Expression levels are coded as per the adjacent keys. **(E)** UMAP of the E17/18 scRNAseq dataset in Figures 2A-C, showing only the clusters containing RP/NSCs and IPs, overlaid for expression of the dorsolateral mRNA *Veph1* versus the dorsomedial mRNA *Slit2*. Expression levels are color-coded as per the adjacent keys. **(F)** Spatial plot of the same ROI as in Figure 5C, overlaid for expression of *Ano1* mRNA, which is enriched at the lateral cortex/GE boundary. Expression levels are color-coded as per the adjacent key. **(G)** UMAPs as in Figure 5I showing thresholded expression of the TAP marker genes *Egfr* (left panel) and *Gsx2* (right panel). Expression thresholds are indicated above each UMAP. Note that *Egfr* is also expressed by OPCs and *Gsx2* by septal and GE IP/RPs. **Abbreviations:** Excit., excitatory. GE, ganglionic eminence. IPs, intermediate progenitors. NSCs, neural stem cells. OPCs, oligodendrocyte precursor cells. Prolif., proliferative. RPs, radial precursors. TAPs, transit-amplifying precursors.

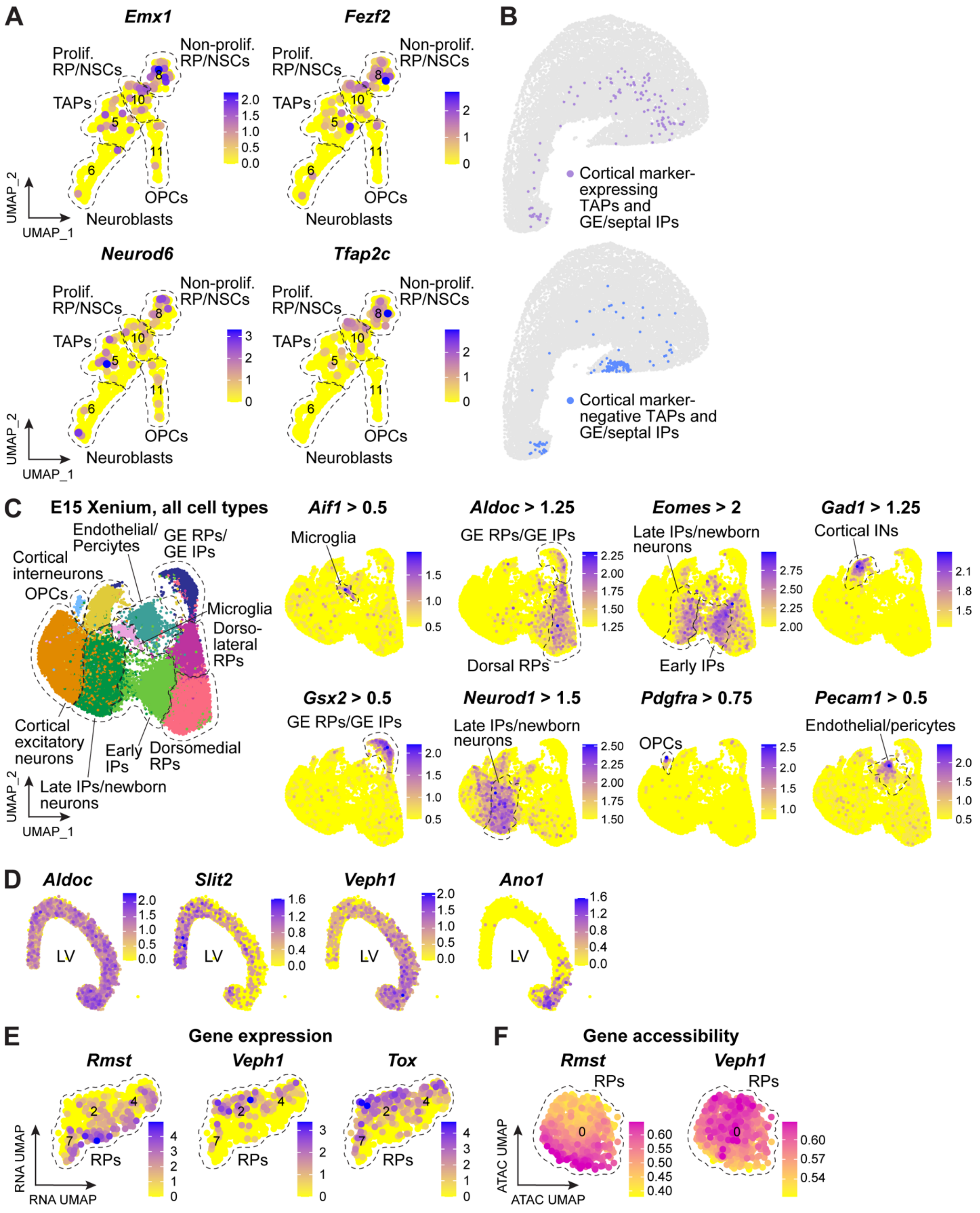


**Figure S6.** ***Definition of cortical TAPs and analysis of E15 Xenium single-cell spatial transcriptomic data*****. (Related to Figure 6). (A)** UMAPs of the E17/18 scRNA-seq V-SVZ cell lineage dataset in Figures 2A-C, showing only the clusters containing RP/NSCs, TAPs, OPCs,

and neuroblasts, overlaid for expression of the cortical mRNAs *Emx1*, *Fezf2*, *Neurod6*, and *Tfap2c*. Expression levels are color-coded as per the adjacent keys and cell types are annotated (indicated by hatched lines). **(B)** Spatial plots of the same representative E18 ROI as in Figure 5C, showing the location of TAPs and GE/septal IPs that do (top panel) or do not (bottom panel) express the cortical marker mRNAs *Emx1*, *Tfap2c*, *Fezf2*, or *Neurod6*. **(C)** UMAPs of the E15 Xenium dataset as in Figure 6F (and shown here on the left) overlaid for thresholded expression of marker genes, including *Aif1* for microglia, *Aldoc* for RP/NSCs, *Gsx2* for GE/septal RPs, *Eomes* for IPs, *Gad1* for inhibitory neuroblasts and neurons, *Neurod1* for late IPs/cortical excitatory neurons, *Pdgfra* for OPCs, and *Pecam1* for endothelial cells. Expression thresholds are indicated above each UMAP. **(D)** Spatial plots of the E15 RPs as defined in Figure 6M, overlaid for expression of mRNAs enriched (C) in all RPs (*Aldoc*), in dorsolateral (*Veph1*) versus dorsomedial (*Slit2*) RPs, or at the lateral cortical/GE boundary (*Ano1*). Expression levels are coded as per the adjacent keys. **(E)** UMAP of the E14 multiomics dataset clustered on the transcriptional data as in Figure 3A, showing only the RP clusters (2, 4, and 7), overlaid for expression of the dorsolateral RP mRNAs *Veph1* and *Tox* versus the dorsomedial RP mRNA *Rmst*. Expression levels are color-coded as per the adjacent keys. **(F)** UMAP of the E14 multiomics dataset clustered on the snATAC-seq data as in Figure 3B, showing only the RP cluster (0), overlaid for chromatin accessibility of the dorsolateral RP gene *Veph1* versus the dorsomedial RP gene *Rmst*. Accessibility is color-coded as per the adjacent keys.
**Abbreviations:** GE, ganglionic eminence. IPs, intermediate progenitors. NSCs, neural stem cells. OPCs, oligodendrocyte precursor cells. Prolif., proliferating. RPs, radial precursors. TAPs, transit-amplifying precursors.

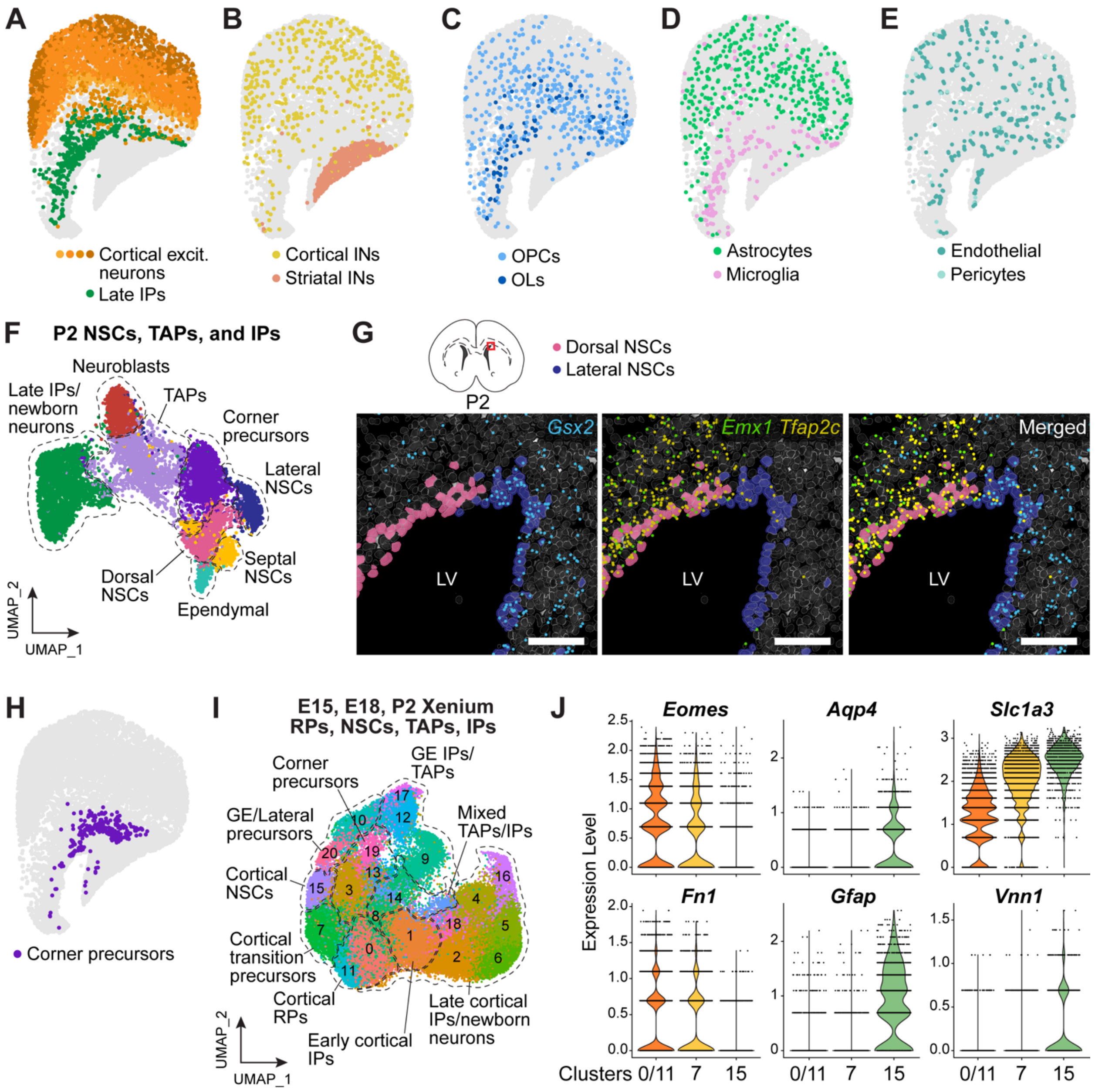


**Figure S7.** ***Analysis of P2 Xenium spatial transcriptomic data and definition of cortical RPs, transition precursors and NSCs in the Xenium datasets.*** **(Related to Figure 7). (A-E)** Spatial plots of the same representative P2 ROI as in Figure 7B, showing cell types plotted individually or in combination, and colored as per the adjacent legends. Shown are (A) late IPs/newborn neurons and more mature cortical excitatory neurons, with different colors denoting transcriptionally- and spatially-distinct groups, (B) cortical and striatal interneurons, (C) OPCs and oligodendrocytes, (D) astrocytes and microglia, and (E) endothelial cells and pericytes. Each dot represents an individual cell. **(F)** UMAP of P2 Xenium neural precursor transcriptomes that were subsetted from the dataset shown in Figure 7A and reanalyzed. Shown are the different cell types, annotated, colored and outlined. **(G)** High-resolution Xenium Explorer images of the dorsolateral V-SVZ (as in the schematic) showing expression of the GE marker mRNA *Gsx2* (blue dots) and the cortical marker mRNAs *Emx1* (green dots) and *Tfap2c* (yellow dots). Also shown are dorsal cortical NSCs (pink cells) and lateral NSCs (dark blue cells). **(H)** Spatial plot of the same representative P2 ROI as in panels A-E and Figure 7B, showing corner precursors in purple. **(I)** UMAP of precursor cell transcriptomes from the E15, E18, and P2 Xenium datasets that were merged together and reanalyzed. Shown are color-coded transcriptionally-distinct clusters, annotated for cell types (indicated by hatched lines), as defined based on marker gene expression, location, and predominant timepoint of origin. **(J)** Violin plots showing normalized expression of the RP-enriched mRNAs *Eomes* and *Fn1*, and the NSC-enriched mRNAs *Aqp4*, *Gfap*, *Slc1a3*, and *Vnn1* in the transcriptionally-distinct RP-like precursors, transition precursors and NSC-like precursors shown in (I) and Figure 7I. Each dot represents expression levels in an individual cell. **Scale bars,** 50 μm. **Abbreviations:** Excit., excitatory. GE, ganglionic eminence. INs, interneurons. IPs, intermediate progenitors. NSCs, neural stem cells. OLs, oligodendrocytes. OPCs, oligodendrocyte precursor cells. TAPs, transit-amplifying precursors.
